# Structural basis for distinct inflammasome complex assembly by human NLRP1 and CARD8

**DOI:** 10.1101/2020.06.17.156307

**Authors:** Gong Qin, Kim Robinson, Xu Chenrui, Zhang Jiawen, Boo Zhao Zhi, Daniel Eng Thiam Teo, Zhang Yaming, John Soon Yew Lim, Goh Wah Ing, Graham Wright, Franklin L. Zhong, Wu Bin, Bruno Reversade

## Abstract

Nod-like receptor (NLR) proteins activate pyroptotic cell death and IL-1 driven inflammation by assembling and activating the inflammasome complex. Closely related NLR proteins, NLRP1 and CARD8 undergo unique auto-proteolysis-dependent activation and are implicated in auto-inflammatory diseases; however, the molecular mechanisms of activation are not understood. Here we report the structural basis of how the activating domains (FIIND^UPA^-CARD) of NLRP1 and CARD8 self-oligomerize to trigger the assembly of distinct inflammasome complexes. Recombinant FIIND^UPA^-CARD of NLRP1 forms a two-layered filament, with an inner core composed of oligomerized CARD domains and the outer layer consisting of FIIND^UPA^ rings. Biochemically, oligomerized NLRP1-CARD is sufficient to drive ASC speck formation in cultured human cells via filament formation-a process that is greatly enhanced by NLRP1-FIIND^UPA^, which forms ring-like oligomers *in vitro*. In addition, we report the cryo-EM structures of NLRP1-CARD and CARD8-CARD filaments at 3.7 Å, which uncovers unique structural features that enable NLRP1 and CARD8 to discriminate between ASC and pro-caspase-1. In summary, our findings provide unique structural insight into the mechanisms of activation for human NLRP1 and CARD8, uncovering an unexpected level of specificity in inflammasome signaling mediated by heterotypic CARD domain interactions.

## Introduction

Nod-like receptor (NLR) proteins are cytosolic sensors and activators of the innate immune response. Many NLR proteins nucleate the ‘inflammasome’-a large macromolecular complex that triggers inflammatory cell death and cytokine secretion-in response to microbial or danger-associated ligands. Like other supramolecular assemblies of the human innate immune system ^1,2^, the inflammasome complex relies on homotypic and heterotypic interactions between the death domains (DDs) of NLRs, adaptor protein ASC and effector caspases to initiate, amplify and propagate pyroptotic signaling ^3,4^. Inflammasome sensor NLRs can be grouped into two categories based on their DDs: PYRIN domain (PYD) and caspase activation and recruitment domain (CARD) ^5–7^. Recently we and others have expanded the repertoire of CARD-based human NLRs to include NLRP1 and CARD8 ^8–11^, both of which are implicated in human Mendelian auto-inflammatory diseases (Fig. 1A) ^9,12–14^. NLRP1 and CARD8 both harbor a poorly defined domain known as FIIND, whose auto-proteolytic activity is required for inflammasome activation ^15,16^. The requirement of the adaptor protein ASC has also been a matter of debate for CARD-containing sensor proteins such as NLRP1 and NLRC4 ^17^. Very recently, human CARD8, which is absent from rodents, was found to bypass ASC altogether and activate caspase-1 directly ^18^.

**Fig. 1.**
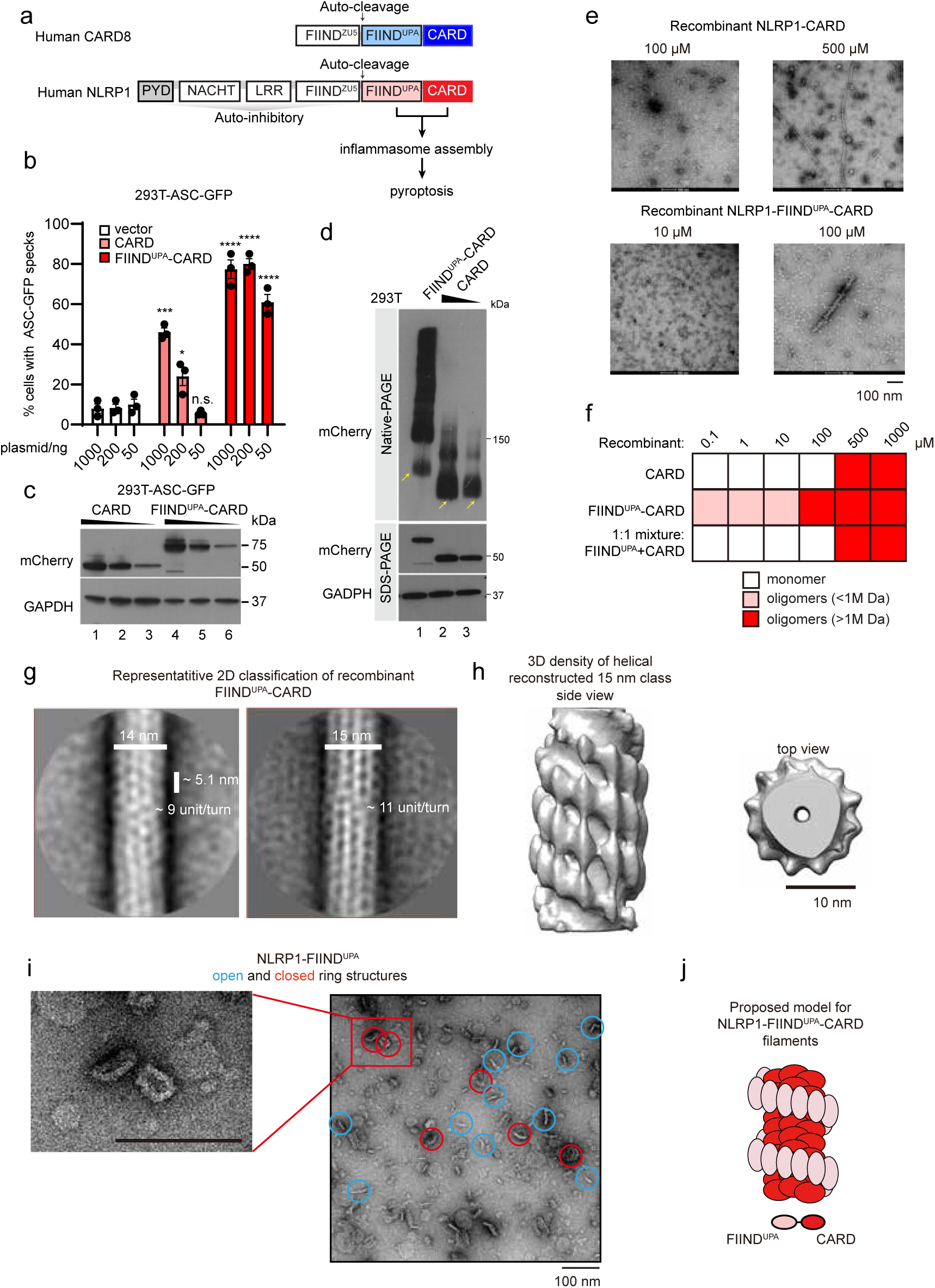
Human NLRP1 and CARD8 nucleate distinct inflammasome complexes. a. Domain organizations of human NLRP1, CARD8. Death domains (PYD and CARD) are highlighted in blue or red. b. Percentage of ASC-GFP specks induced by different amounts of mCherry-tagged NLRP1-FIIND^UPA^-CARD (red) or mCherry-tagged NLRP1-CARD (light red) in HEK293T-ASC-GFP cells. Cells were fixed 24 hours after transfection for fluorescence microscopy. P-value was calculated with One-way ANOVA, n=3 biological replicates. c. Corresponding Western blot of mCherry-tagged NLRP1 constructs in Fig. 1B. d. Blue-Native PAGE blot of mCherry-tagged NLRP1-FIIND^UPA^-CARD and mCherry-tagged NLRP1-CARD. Yellow arrows, monomeric species. e. Representative negative stain EM images recombinant NLRP1-FIIND^UPA^-CARD and NLRP1-CARD at various concentrations. f. A summary of oligomer formation of various NLRP1 constructs at various concentrations. g. Two of the best 2D class images with clearly distinguishable features of recombinant NLRP1-FIIND^UPA^-CARD filamentous complex, from negative stain EM analysis. h. A side view and a top view of the 3D density model at estimated 22 Å resolution (after 3D refinement using RELION) generated based on particles extracted from the 15 nm 2D class. i. Negative EM images of SUMO-tagged NLRP1-FIIND^UPA^. The ring-complexes are roughly 20 nm in size. j. Proposed model for NLRP1 FIIND^UPA^-CARD oligomers.

In this study, we investigate the structural basis of how the auto-proteolysis-dependent, activating domains of human NLRP1 and CARD8 (FIIND^UPA^-CARD) trigger inflammasome assembly and activation. We demonstrate that FIIND^UPA^ and CARD domains play non-redundant roles in NLRP1 activation: NLRP1-CARD is sufficient to form filaments *in vitro*, trigger ASC speck formation and activate pyroptosis in human cells, albeit with limited efficiency. The FIIND^UPA^ domain of NLRP1 does not interact with ASC directly, but dramatically lowers the threshold of NLRP1-CARD oligomerization and filament formation. Mechanistically, a stand-alone FIIND^UPA^ forms ring-like oligomers while recombinant FIIND^UPA^-CARD forms a two layered filament, with oligomerized CARD as the inner core around which a FIIND^UPA^ ring spirals. By solving the cryo-EM structures NLRP1- and CARD8-CARD filaments at a resolution of 3.7 Å, we map the precise structural motifs that mediate specific heterotypic interactions between the CARD domains of NLRP1, CARD8, ASC and CASP1. This analysis provides a comprehensive structural explanation as to why activated NLRP1 avidly triggers ASC speck formation, while CARD8 directly activates pro-caspase-1. Taken together, our findings provide fresh structural insight on human inflammasome assembly and reveal an unexpected level of specificity in inflammasome signaling conferred by the intrinsic structural properties of the CARD domain.

## Results

### NLRP1-FIIND^UPA^ domain facilitates NLRP1-CARD oligomerization

Previously we and others have demonstrated that C-terminal auto-cleavage fragments of NLRP1 and CARD8 (NLRP1-FIIND^UPA^-CARD and CARD8-FIIND^UPA^-CARD) are sufficient to drive pyroptosis in cultured human cells ^9,11,19^ (Fig. 1a). To further map the exact roles played by FIIND^UPA^, we utilized the widely used ‘ASC-speck’ assay in HEK239T cells expressing GFP-tagged full-length ASC (293T-ASC-GFP). Consistent with previous results, we found that overexpressed NLRP1-CARD can be sufficient to induce ASC-GFP specks, but does so much less readily without the FIIND^UPA^ domain. This difference is especially notable at lower levels of protein expression (Fig. 1b, 50 ng; Fig. 1c, lane 3 and 6). As NLRP1 activation is strictly correlated with its self-oligomerization^9^, we examined the effect of the FIIND^UPA^ domain on NLRP1-CARD oligomerization by native PAGE. At limiting protein expression levels, ∼90% of NLRP1-CARD exists in the monomeric form (Fig. 1d, lane 2 and 3), with the rest as higher-molecular weight oligomers, in agreement with its low but detectable activation capacity. In contrast, FIIND^UPA^-CARD, at comparable expression levels, exists mostly as high-molecular weight oligomeric species (Fig. 1d, lane 1 vs. 2 and 3). Taken together, these results suggest that while NLRP1-CARD is sufficient to induce ASC specks in its oligomerized form, the presence of FIIND^UPA^ domain strongly enhances its activation by lowering the threshold of self-oligomerization.

Next we examined purified recombinant NLRP1-CARD, FIIND^UPA^-CARD, and a 1:1 molar mixture of FIIND^UPA^ + CARD using negative stain electron microscopy (EM). In agreement with the native PAGE results in 293T cells, NLRP1-FIIND^UPA^ effectively lowered the concentration requirement for CARD oligomerization by at least 50 fold (Fig. 1e and 1f), but only when expressed together with NLRP1-CARD as a single polypeptide (Fig. 1f). Mixing NLRP1-FIIND^UPA^ with CARD had no effect (Fig. 1f, bottom row). NLRP1-CARD, in isolation or incubated with FIIND^UPA^ as a separate polypeptide, was unable to form ordered oligomers at concentrations <∼200 μM, but did appear as long ordered oligomers at concentrations >500 μM (Fig. 1e, right, Fig. 1f). In keeping with the cellular experiments, these results suggest that in the absence of FIIND^UPA^, NLRP1-CARD has a limited but detectable propensity for self-oligomerization and activation. The FIIND^UPA^ domain, akin to a ‘built-in’ catalyst, acts as potent facilitator for NLRP1-CARD oligomerization and dramatically lowers its threshold for activation

To seek direct structural evidence for this model, we purified the following NLRP1 fragments: FIIND^UPA^-CARD, full length FIIND-CARD (FIIND^fl^-CARD), FIIND^UPA^ and CARD alone. Both HEK293T cell-derived and bacteria-derived FIIND^UPA^-CARD formed long heterogenous filaments approximately 14-15 nm in diameter (Fig. S1a-b), but not FIIND^UPA^ alone (Fig. S1c), or FIIND^fl^-CARD (Fig. S1d). In agreement with its detectable oligomerization capacity in 293T cells, recombinant NLRP1-CARD also formed filaments, which were visibly thinner than those formed by FIIND^UPA^-CARD (∼ 8 nm in diameter) (Fig. S1b). To elucidate the relative arrangement of FIIND^UPA^ and CARD within the FIIND^UPA^-CARD structure, we examined the topology of recombinant FIIND^UPA^-CARD filaments using cryo-EM. Multiple 2D classes with distinct rotational symmetries were present with various diameters and helical settings (Fig. 1g). 3D projection of the most stable 15 nm 2D class yielded a helical filament with 11-12 identical subunits per turn and ∼5.6 nm per helical rise. Notably, when examined using differing contour levels, this FIIND^UPA^-CARD filament class was found to consist of a denser inner core filament with a diameter of ∼8 nm (Fig. 1h). Surrounding this inner core is a discrete outer layer that extends to 15 nm from the center, decorated with distinct ‘blobs’ of protruding density (Fig. 1h). To the best of our knowledge, this is the first time such a ‘two-layer’ architecture has been observed for a FIIND-containing inflammasome sensor. It is remarkably reminiscent of activated NLRC4 and NLRP6 ^20,21^, where the nucleotide-binding domain (NBD) pre-oligomerizes to bring the CARD or PYRIN domains into close proximity. Furthermore, the observed dimensions of the inner core filament is consistent with those of the recombinant filaments composed of CARD alone (Fig.S1b, 8 nm in diameter, see below). Therefore, we propose that the FIIND^UPA^-CARD oligomers, which represent the activated form of NLRP1, consist of an inner helical filament of oligomerized CARDs surrounded by a FIIND^UPA^ spiral facing outward (Fig. 1i). As further evidence for this model, recombinant SUMO-tagged NLRP1-FIIND^UPA^ spontaneously formed ring-shaped oligomers of heterogeneous stoichiometries *in vitro*. These closed ‘rings’ had a minimum diameter of ∼17 nm (Fig. 1i, red circles). Accommodating the SUMO tag, this is consistent with the dimensions of the outer periphery of the NLRP1-FIIND^UPA^-CARD filament (Fig. 1i and 1j). Similar biochemical analysis of the FIIND^UPA^-CARD and FIIND^UPA^ domains from CARD8 revealed that CARD8-FIIND^UPA^ most likely functions in a similar way as that of NLRP1. Recombinant CARD8-FIIND^UPA^ also formed ring-like oligomers, although the oligomer architecture appears to be more heterogeneous in nature (Fig. S2a-c). Based on these findings, we propose a stepwise model for NLRP1 and CARD8 activation. Once triggered, NLRP1-FIIND^UPA^ and CARD-FIIND^UPA^ pre-oligomerizes to bring the respective CARD domains in close proximity, thereby increasing its potential for filament formation. Once formed, the oligomerized CARDs are sufficient to activate downstream inflammasome signaling. This model is also likely to be valid for NLRP1/CARD8 homologues from other species, as the arrangement of the FIIND and CARD domains are conserved across evolution^22, 23^.

### Cryo-EM structures of NLRP1- and CARD8-CARD filaments: the inner core of the FIIND^UPA^-CARD filaments

It was very recently reported that human NLRP1 and CARD8, long thought to be functional paralogues, can give rise to distinct signaling outcomes: while NLRP1 avidly triggers ASC speck formation, CARD8 can directly activate pro-caspase-1 without the need for ASC in HEK293T cells^18^. The mechanistic basis for this disparity is currently unclear. Considering our finding demonstrating that NLRP1-CARD, once oligomerized, is sufficient to activate downstream inflammasome signaling (Fig. 1b), we hypothesized that this difference could be due to the intrinsic structural differences between NLRP1-CARD and CARD8-CARD oligomers. To test this, we solved the structures of recombinant NLRP1-CARD and CARD8-CARD filaments using cryo-EM, both at 3.7 Å, Gold standard FSC = 0.143, without masking (Fig. 2a-i, Fig. S3a and Supplementary Table 1). NLRP1-CARD filaments are relatively flexible and adopt multiple helical organizations *in vitro*, as evidenced by heterogeneous 2D and 3D classes observed during structural analysis using RELION 3.0 (Fig. S3b) ^24^. One out of four observed 3D class conformations of NLRP1-CARD filaments was determined to be suitable for high-resolution structural determination ^25^ (Fig. S3c, Fig. 2b-e). This conformation was subsequently verified to be a signaling-competent conformation based on functional experiments using 293T-ASC-GFP cells (see below). Importantly, the resolved CARD filament density fits snugly into the lower resolution model of NLRP1-FIIND^UPA^-CARD filament based on negative stain EM, matching the shape and dimensions of the inner layer almost perfectly (Fig. 2e). Similar to NLRP1-CARD, recombinant CARD8-CARD also formed ∼8 nm-wide filaments, but these filaments are more rigid and adopt a single conformation (Fig. 2f-h, and Fig. S4a-b). The final polished densities demonstrated clear α-helix pitch and supported the unambiguous assignment of all bulky side chains (Fig. 2d and 2i, Fig. S4c-d, Fig. S4a-d) in both filaments.

**Fig. 2.**
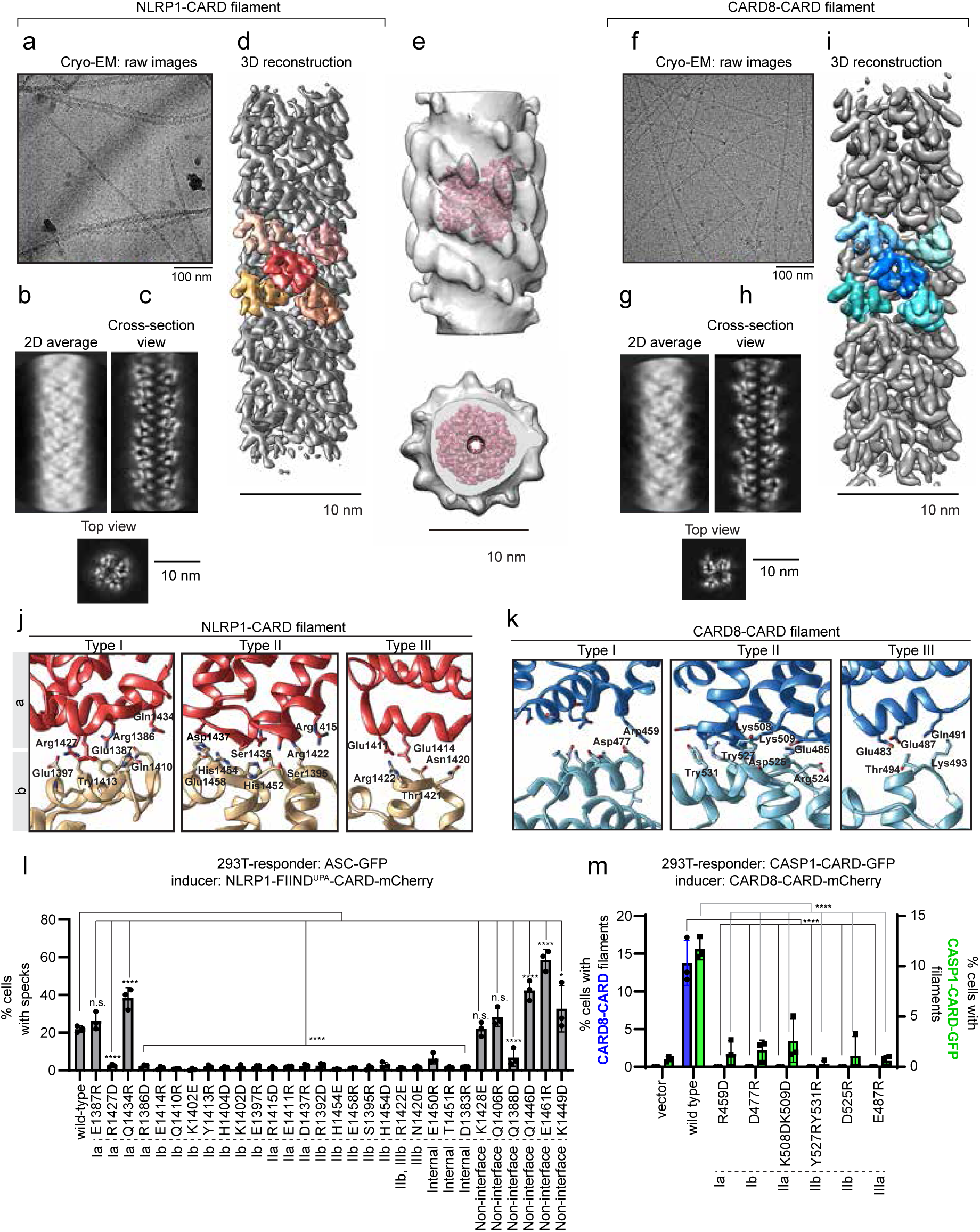
Cryo-EM structures of NLRP1-CARD and CARD8-CARD filaments. a. Cryo-EM images of recombinant NLRP1-CARD filaments. b. 2D average of NLRP1-CARD filaments. c. Cross-section and top view of NLRP1-CARD filaments. d. Helical reconstructed NLRP1-CARD complex density. Adjacent NLRP1-CARD monomers are colored. e. Atomic resolution NLRP1-CARD filament density fit into the inner layer density of NLRP1-FIIND^UPA^-CARD shown in Fig. 1h-1i. f. Cryo-EM raw images of recombinant CARD8-CARD filaments. g. 2D average of CARD8-CARD filaments. h. Cross-section and top view of CARD8-CARD filaments. i. Helical reconstructed CARD8-CARD complex density. Adjacent CARD8-CARD monomers are colored. j. Type I-III interfaces in NLRP1-CARD filaments. k. Type I-III interfaces in CARD8-CARD filaments. l. Mutagenesis of critical NLRP1-CARD interface residues and their effects on NLRP1-FIIND^UPA^-CARD-mCherry to induce ASC-GFP specks in 293T cells. P-value was calculated with One-way ANOVA, n=3 biological replicates. m. Mutagenesis of critical CARD8-CARD interface residues and their effects on CARD8-CARD-mCherry filament formation (blue) and CASP1-CARD-GFP oligomerization in 293T cells (green).

NLRP1-CARD and CARD8-CARD filaments display similar helical parameters (5.36 Å and -100.8 degree for NLRP1, and 5.409 Å and -99.16 degree for CARD8), suggesting an overall similar mode of assembly from the respective monomers (Fig. 2d, 2i). Both filaments also have clearly demarcated Type I-III interfaces that are characteristic of oligomerized Death Domains (DD) (Fig. 2j-k, Fig. S4c-d) ^2,20,26,27^. As compared to other DD complexes the Type II interfaces in NLRP1- and CARD8-CARD filaments display stronger and more prominent electron density than Type I and Type III interfaces (Fig. 2j-k). This is particularly notable in CARD8-CARD filaments, where Type Ia and Ib surface are more separated from each other (Fig. 2j-k). To validate these structures functionally, predicted Type I-III interface residues were mutated by site-directed mutagenesis. For both filaments, missense mutations of Type I-III interface residues abrogated filament assembly as well as their ability to nucleate downstream ASC-GFP or CASP1-CARD-GFP specks/filaments (Fig. 2l-m), whereas non-interface surface-exposed residues have less effect. These results further prove that CARD oligomerization is a prerequisite for NLRP1 and CARD8-driven inflammasome activation. A notable exception is the Type Ia interface of NLRP1, where two of the four mutations, E1387R and Q1434R, did not affect ASC-GFP speck formation in HEK293T cells despite partially reducing NLRP1-CARD filament assembly *in vitro* (Fig. 2l, Fig. S3d). We postulate that the Type Ia (bottom-facing) interface of NLRP1 is predominantly involved in NLRP1-CARD self-oligomerization, rather than NLRP1-ASC interaction.

### NLRP1 and CARD8 CARD oligomers have distinct surface charge distributions despite similar helical symmetry

Despite near-identical helical parameters, NLRP1-CARD and CARD8-CARD filaments have remarkably different surface charge distributions (Fig. 3a-f). NLRP1-CARD filaments demonstrate a balanced charge distribution on its axial surface, similar to previously published DD complexes (Fig. 3a) ^2,4,28^. However, in CARD8-CARD filaments, the positively charged patches on the monomer are mostly buried inside the filamentous oligomer, exposing a predominantly negative surface (Fig. 3b). The density from the Type I interface, which plays a prominent role in supporting the oligomerization of other CARD helical complexes, does not seem to play a major role in the observed surface density in either complex. Instead, the surface charge distribution is largely due to unusually strong Type II interface densities in both NLRP1-CARD and CARD8-CARD filaments. In the NLRP1-CARD complex, two positively charged histidine residues (His1454 and His1452) extend from the Type IIb surface and interact with negatively charged residues, Asp1437 and Ser1435 on the Type IIa surface (Fig. 3c). In contrast, the CARD8 Type IIb surface is predominantly negatively charged (Tyr531, Tyr527 and Asp525) whereas the Type IIa surface is positive (Lys508, Lys509 and Leu512) (Fig. 3d). This distinct arrangements of the Type II interfaces are a direct result of the subtle differences (2 degree in helical rotation and ∼15 degree of monomer rotational position) in their helical symmetries, and therefore not easily predicted from the amino acid sequence differences between NLRP1 and CARD8. Interestingly, the top surface of NLRP1-CARD closely resembles the charge distributions of ASC-CARD oligomer top surface (ASC-CARD filament solved in-house, PDB-6K99, EMD-9947, Fig. 3e, Fig. 3g, left), and CARD8 top surface is more similar to CASP1-CARD top surface (Fig. 3f, Fig. 3h, left). Hence, NLRP1-CARD top surface is predicted to preferentially interact with the bottom surface of ASC-CARD (compare Fig. 3e and 3g, right), whereas the CARD8-CARD top surface is expected to preferentially interact with the CASP1-CARD bottom surface (compare Fig. 3f and 3h, right). These structural findings offer a potential explanation for the differing affinity and requirement for ASC between NLRP1 and CARD8, which was recently inferred on the basis of cellular experiments.

**Fig. 3.**
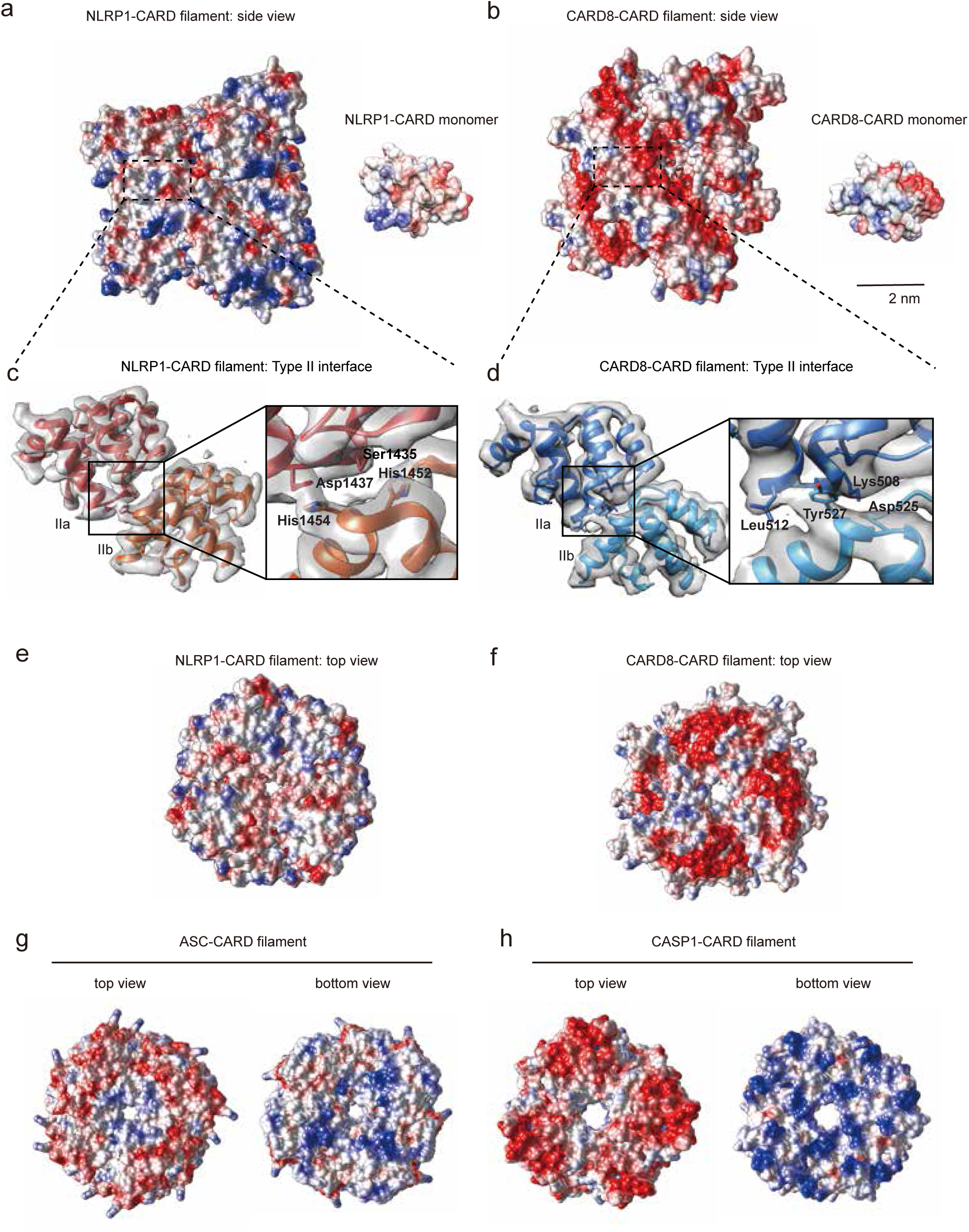
Oligomeric interfaces on inflammasome CARD domains. a. Left panel: side-view of electrostatic potential distribution of a segment of NLRP1-CARD filament (red, -10 *C*/m^2^; blue, 10 *C*/m^2^; generated in Chimera). Right panel: electrostatic potential map of a NLRP1-CARD monomer in the same orientation as the filamentous oligomer. b. Left panel: side-view of electrostatic potential distribution of a segment of CARD8-CARD filament (red, -10 *C*/m^2^; blue, 10 *C*/m^2^). Right panel: electrostatic potential map of a CARD8-CARD monomer in the same orientation. c. Atomic model cartoon and the directly observed density map of a NLRP1-CARD dimer within the filamentous complex, centered at the Type II junction interface. The four residues, His1452-His1454 on the Type IIb interface and the Ser1435-Asp1437 on the Type IIa interface were highlighted and shown in the ‘stick’ mode. d. Atomic model cartoon and the directly observed density map of CARD8-CARD dimer in the filament, centered at the Type II junction interface. The four residues, Asp525-Tyr527 on the Type IIb interface and the Lys508-Leu512 on the Type IIa interface were highlighted and shown in the ‘stick’ mode. e. Electrostatic potential map of the top view (Type IIb interface, highlighted in dashed area) of the NLRP1-CARD filament. f. Electrostatic potential map of the top view (Type IIb interface, highlighted in dashed area) of CARD8-CARD filament. g. Electrostatic potential map of the top and bottom views of the ASC-CARD filament. h. Electrostatic potential map of the top and bottom views of the CASP1-CARD filament.

### Structural basis for the distinct downstream CARD preference between NLRP1 and CARD8

To identify the precise structural determinants that are responsible for ASC vs. pro-caspase-1 binding, we focused our attention on the Type II surfaces of the respective CARD oligomers. This surface plays a prominent role in the observed differences in surface charge distribution between NLRP1-CARD and CARD8-CARD filaments (Fig. 2j-k, Fig. 3a-f). We hypothesized that the Type II junctions are not only important for homotypic assembly between identical CARD subunits, but also heterotypic interactions between ‘sensor’ CARDs (NLRP1 and CARD8) and downstream effector CARDs (ASC and CASP1). To test this, we first identified the Type II interfaces in published ASC-CARD and CASP1-CARD filament structures (Fig. S5a-e, Supplement Table 1). In both structures, we noted a similar internal density bridge on the Type II interface as what was observed in NLRP1-CARD and CARD8-CARD complexes (compare Fig. 4a-b to Fig. 3c-d), which had not been highlighted by previous analyses ^20,27,29–35^. In ASC-CARD filaments, this bridge consists of two residues Gln185 and Tyr187, which extend from the Type IIb surface and interact with Ala168 and Asn170 on the Type IIa surface (Fig. 4a). In CASP1-CARD filaments, the Type II interface bridge employs a distinct IIa surface composed of Lys64 and Gln67, which interacts with Type IIb residues, Asp80 and Tyr82 (Fig. 4b). This ‘bridge-like’ feature across the Type II interface becomes even more apparent when we refined the existing CASP1-CARD model (Fig. S5c). Hence a strong inter-subunit Type II density bridge seems to be a common feature amongst inflammasome CARD complexes.

**Fig. 4.**
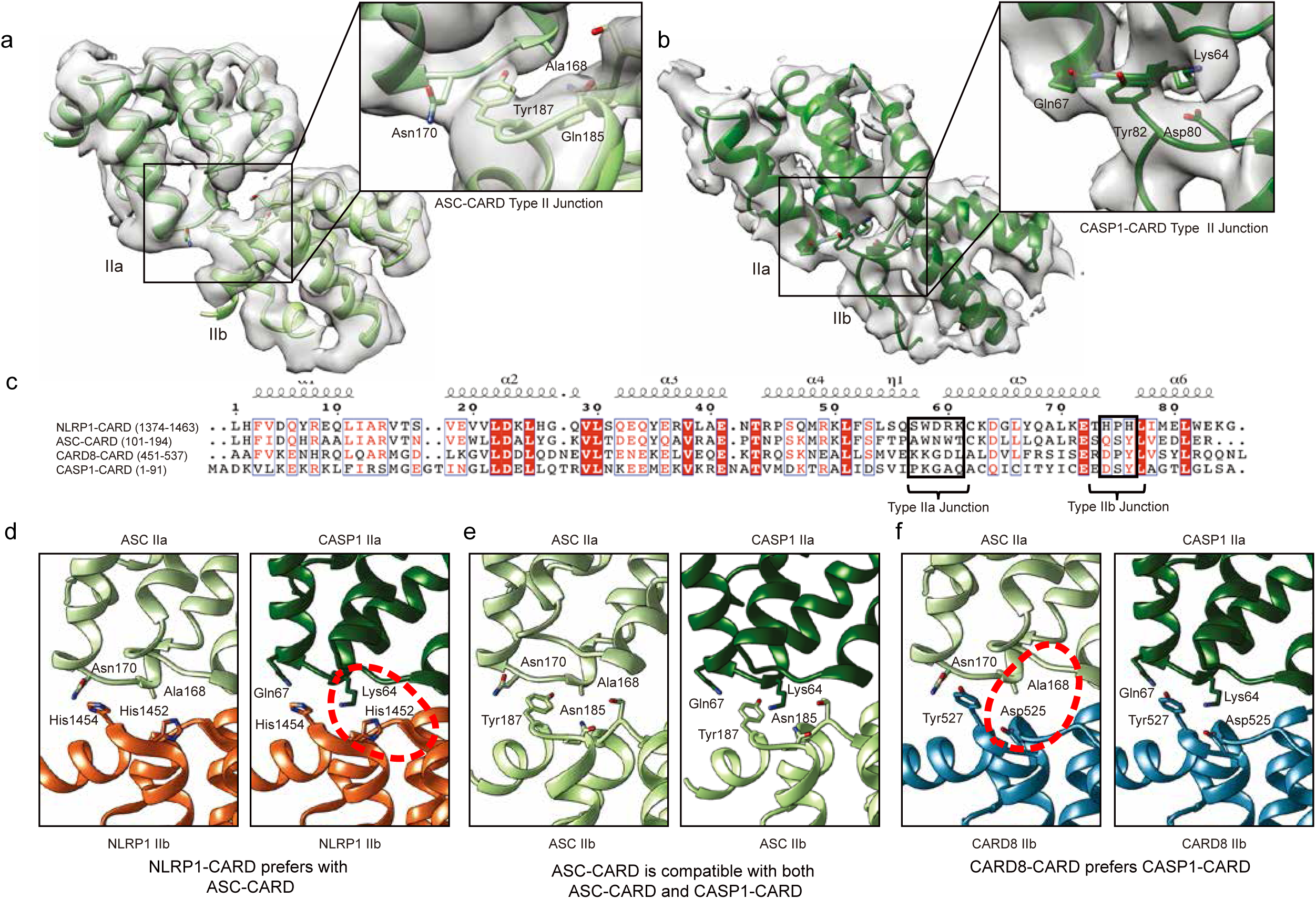
Type IIa-b interface influences human inflammasome complexes. a. Type II interface in ASC-CARD filaments (PDB-6K99, EMD-9947). b. Type II interface in CASP1-CARD filaments after refitting. See Fig.S5c for a comparison between the refitted model and the original model, PDB-5FNA and EMD-3241). c. Sequence alignment of human NLRP1-, ASC-, CARD8- and CASP1-CARD. Important residues within the Type II interfaces are highlighted in black box. d. Structural compatibility of human NLRP1-CARD with ASC-CARD or CASP1-CARD. Potential clashes are highlighted in red. e. Structural compatibility of human ASC-CARD with itself or CASP1-CARD. f. Structural compatibility of human CARD-CARD with ASC-CARD or CASP1-CARD. Potential clashes are highlighted in red.

Using the refined assignment of Type II interface residues, we modelled all the pairwise unidirectional CARD-CARD interactions amongst NLRP1, CARD8, ASC and CASP1 (Fig. 4c-f), based on our in-house built filament models (PDB-6K7V, 6K9F, 6K99) and re-refined CASP1-CARD oligomer (locally refined PDB-5FNA using our CARD8-CARD structure as a reference). A remarkable level of specificity becomes apparent, as certain interactions are favored while others show patent incompatibility. NLRP1-CARD and ASC-CARD is expected to be compatible, as the prominent HPH (a.a. 1452-1454) motif at the NLRP1 Type IIb interface can form ionic interactions with the ASC IIb residues AWN (a.a. 168-170) (Fig. 4c, Fig. 4d, left panel). In contrast, the same NLRP1 HPH motif clashes with the CASP1 Type IIa surface residues, KGAQ (a.a. 64-67) (Fig. 4c, Fig. 4d, right panel, circle). In other words, NLRP1-CARD uses the HPH motif on its Type IIa surface to preferentially engage ASC-CARD and disfavor CASP1-CARD (Fig. 4c-d). On the other hand, CARD8-CARD employs a DPY (a.a. 525-527) motif on its Type IIb surface. This allows CARD8-CARD to interact strongly with the CASP1-CARD Type IIa surface via ionic interactions between Asp 525 (CARD8) and Lys64 (CASP1) (Fig. 4c, Fig. 4f, right panel). In contrast, the same DPY motif lacks effective interaction with the ASC-Type IIa surface, as the charged Asp 525 in the CARD8 DPY motif cannot pair with Ala 168 on the ASC Type IIa surface (Fig. 4f, left panel). Hence CARD8-CARD is structurally compatible with CASP1-CARD but not ASC-CARD. These results demonstrate that NLRP1-CARD and CARD8-CARD have the intrinsic ability to discriminate between ASC and pro-caspase-1 due to the structural features of their Type IIb motifs.

This analysis can be further extended to explain why ASC is uniquely suited to act as an adaptor and amplifier in inflammasome signaling. To do so, ASC-CARD must self-oligomerize while remaining fully capable of engaging CASP1-CARD. This versatility is endowed by its unique Type IIb motif, QSY (a.a. 185-187), which possesses biochemical properties and charges that are intermediate between NLRP1-CARD and CARD8-CARD. The Type IIb QSY motif in ASC-CARD adopts a conformation that is compatible both with its own Type IIa residues, i.e. a.a. AWN (a.a. 168-170), as well as the CASP1 ‘KGAQ’ motif (Fig. 4e). This is achieved with the help of additional sidechain features. For instance, instead of a prominent negative Asp residue in CARD8 Type IIb surface, ASC-CARD uses a less charged Asn residue to enhance the interaction with Ala168 on its own type IIa surface. In addition, the tyrosine residue (Tyr 187) on ASC-CARD Type IIb demonstrate a more inwards orientation as compared to the equivalent tyrosine residue on CARD8 type IIb surface (Tyr 527), thus allowing it to effectively engage both its own Type IIa as well as that of CASP1-CARD (Fig. 4e).

To further corroborate that signaling specificity is intrinsic to the CARD domains, we performed an *in vitro* oligomerization assay using recombinant fluorescently-labelled CASP1-CARD and ASC-CARD. In agreement with the structural predictions, CARD8-CARD, but not NLRP1-CARD, preferentially stimulated CASP1-CARD oligomerization, as demonstrated by the appearance of slower migrating bands on native gels (Fig. S5f, left panel, lanes 2 and 3 vs. 4 and 5), while the reverse was true for ASC-CARD oligomerization (Fig. S5f, right panel, lanes 2 and 3 vs. lanes 4 and 5). Next, we used structured illumination microscopy (3D-SIM) to directly visualize NLRP1-CARD and CARD8-CARD and their interactions with downstream CARDs (ASC-CARD vs. CASP1-CARD) within cultured human cells. Both mCherry-tagged NLRP1-CARD and CARD8-CARD formed filament bundles within the cytosol (Fig. 5a, red), which intertwined with those formed by GFP-tagged ASC-CARD and CASP1-CARD, respectively (Fig. 5a, green). Given the theoretical resolution of ∼110 nm in width (x-y plane), we postulated that the thinnest filament bundle observed was no more than approximately 6 times the width of an individual filament with the mCherry tag. In many instances, ASC-CARD or CASP1-CARD filaments could be observed as ‘emanating’ from one side of NLRP1-CARD or CARD8-CARD filament bundles (Fig. 5a). This ‘filament-seeding’ phenomenon was not limited to the isolated CARDs, as Talabostat-activated full-length NLRP1 and CARD8 also formed filaments ^9,15,19^ and colocalized with ASC-CARD and CASP-1 CARD filaments, respectively (Fig. S6). Furthermore, Talabostat-activated full-length NLRP1, or its activating fragment NLRP1-FIIND^UPA^-CARD are unable to trigger CASP1-GFP filaments in the absence of ASC, while CARD8 (or CARD8-FIIND^UPA^-CARD) was unable to nucleate ASC-GFP specks in 293T cells (Fig. 5a-b). Taken together, these results provide direct cellular evidence that NLRP1-CARD and CARD8-CARD form filament-like oligomers that differentially engage either ASC or pro-caspase-1, via heterotypic CARD interactions.

**Fig. 5.**
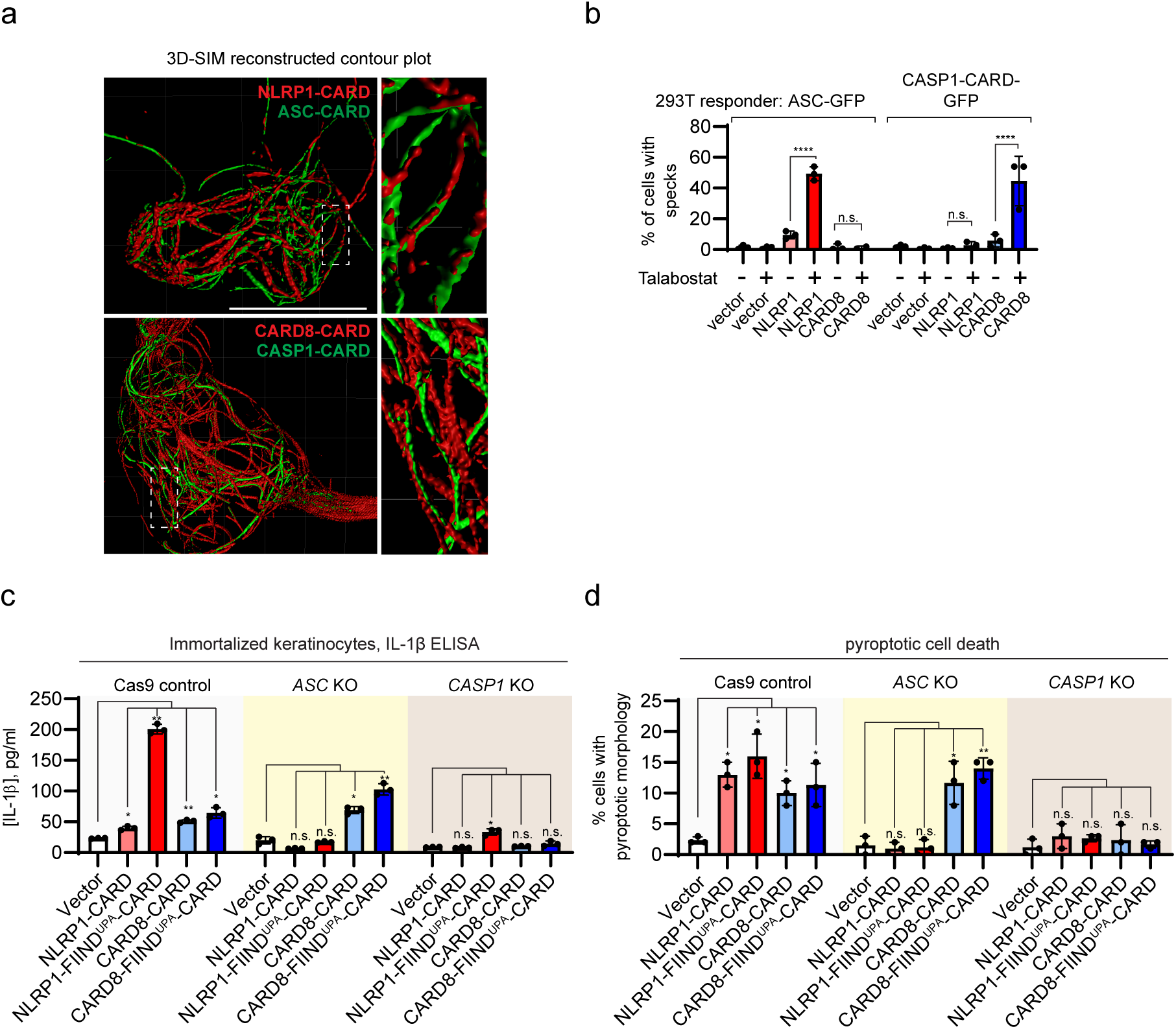
NLRP1 and CARD8 inflammasome assembly in mammalian cells. a. Structured illumination microscopy images of NLRP1-CARD-mCherry and CARD8-CARD-mCherry filaments (red) with ASC-CARD-GFP and CASP1-CARD-GFP filaments (green). White boxes indicate regions magnified in right panels. b. Percentage of ASC-GFP specks or CASP1-CARD-GFP filaments induced by full-length NLRP1 or CARD8 upon Talabostat treatment (3 μM). P-value was calculated with One-way ANOVA, n=3 treatments. c. IL-1β secretion in control, *ASC* KO, and *CASP1* KO keratinocytes transfected with the indicated NLRP1 and CARD8 constructs. P-value was calculated with One-way ANOVA, n=3 transfections. d. The percentage of cells with pyroptotic morphology in control, *ASC* KO, and *CASP1* KO keratinocytes transfected with the indicated NLRP1 and CARD8 constructs. P-value was calculated with One-way ANOVA, n=3 transfections (same as 5c).

We further confirmed that CARD8 is the first human inflammasome sensor that is unable to engage ASC as an amplifier^18^ using immortalized human keratinocytes. In immortalized human keratinocytes, CRISPR/Cas9-mediated deletion of endogenous *ASC* (*PYCARD*) abrogated the ability of overexpressed NLRP1-FIIND^UPA^-CARD fragment or NLRP1-CARD to induce endogenous IL-1β secretion (Fig. 5c, red bars, Fig. S7c-f) and pyroptotic cell death (Fig. 5c-d, red bars, Fig. S7b, S7d). In contrast, CARD8-dependent pyroptosis was not abrogated in *ASC* (*PYCARD*) KO keratinocytes (Fig. 5c-d, blue bars, Fig. S7c, S7e). These results confirm that NLRP1 requires *endogenous* ASC for inflammasome activation in human cells, whereas CARD8 can directly activate caspase-1 independently of ASC.

### NLRC4-CARD can engage either ASC-CARD or CASP1-CARD

Our results thus far demonstrate that subtle differences in the inflammasome CARD domain structures, particularly at the type II junctions, can directly dictate the signaling specificity of the NLR sensor proteins. To extend these results beyond NLRP1 and CARD8 and test the predictive power of our model, we examined a third CARD-containing NLR sensor, NLRC4. To unambiguously assign the Type II interface residues of human NLRC4-CARD, we determined its filament structure at 3.3 Å using cryo-EM (Fig. S5d-e). The final polished structure closely agrees with previously published findings ^20,34^. Interestingly, unlike NLRP1-CARD and CARD8-CARD, the strong density bridge in the Type II interface is absent within NLRC4-CARD filaments (Fig. S8a). The Type IIb tyrosine residue (Tyr78) is folded backward toward the hydrophobic core and does not participate in intermolecular Type II interaction. Instead, two nearby small residues, Asn77 and Gln82, come forward to form a new class of Type II junction (Fig. S8a-b) that is more flexible than that of NLRP1 and CARD8. Molecular docking predicted that NLRC4 Type IIb surface would be structurally compatible with both ASC-CARD and CASP1-CARD. This was validated using the *in vitro* EMSA assay (Fig. S5f, lanes 6-7 in both panels) and in 293T cells. Overexpressed NLRC4-CARD nucleated the formation of both ASC-GFP and CASP1 C285G-GFP specks, as well as ASC-CARD-GFP and CASP1-CARD-GFP filaments at similar levels (Fig. S8c-d). The versatility of human NLRC4-CARD is in agreement with previous studies on murine Nlrc4. In mice, *Asc/Pycard* knockout attenuates, but does not abolish caspase-1-dependent pyroptotic cell death (Dick et al., 2016), suggesting that murine NLRC4, similar to its human homologue can engage both ASC and pro-caspase-1 (Fig. S8e).

## Discussion

Taken together, our cellular, biochemical and structural findings suggest a model for NLRP1- and CARD8-initiated Inflammasome activation (Fig. 6). We propose that FIIND auto-cleavage, which has been shown previously to be required for NLRP1 activation, serves as a necessary ‘licensing’ step to liberate the FIIND^UPA^ domain. Upon ligand-induced activation and relief of auto-inhibition, FIIND^UPA^ self-assembles into ring-like oligomers, bringing adjacent CARD monomers into close proximity. This lowers the threshold CARD oligomerization and allows for further ‘prionioid’-like growth of FIIND^UPA^-CARD filaments (Fig. 6a). Since FIIND auto-cleavage is also required for CARD8 oligomerization, we posit that this mechanism involving FIIND^UPA^-aided CARD oligomerization likely also holds true for the CARD8 inflammasome (Fig. 6b).

**Fig. 6.**
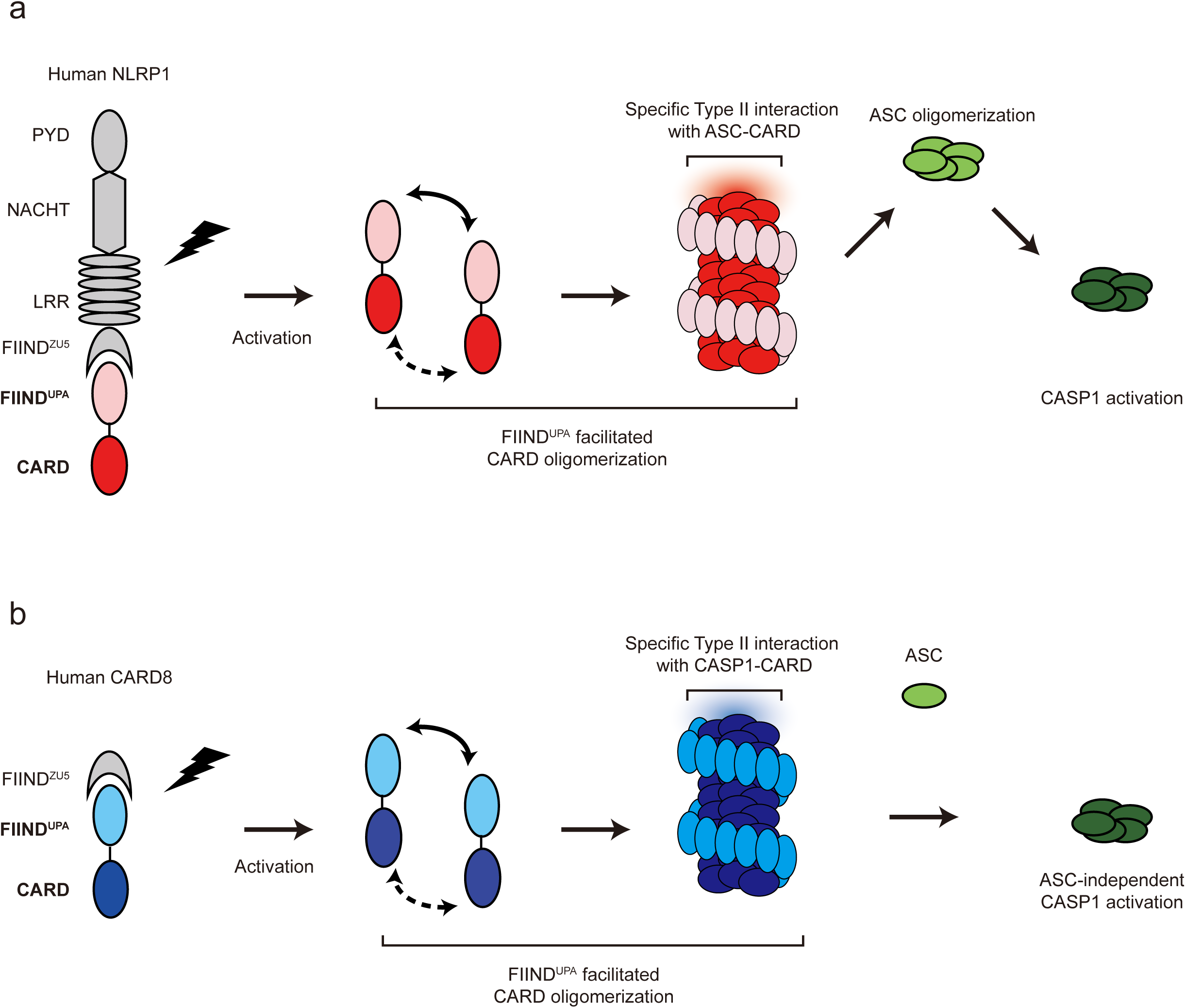
A two-step model for human NLRP1 and CARD8 inflammasome assembly. From left to right, NLRP1 (a) and CARD8 (b) are ‘licensed’ by auto-proteolysis within the FIIND. Upon activation by PAMP or DAMP triggers, the self-inhibition exerted by the N-terminal proteolytic fragment is relieved. FIIND^UPA^ facilitates the oligomerization of CARD, leading to the formation of a two-layered filament complex. Oligomerized NLRP1-CARD and CARD8-CARD differentially engage ASC or CASP1, leading to the assembly of distinct inflammasome complexes.

Since both human NLRP1 and CARD8 employ oligomerized CARDs as a signaling platform to initiate downstream inflammasome signaling, we next sought to solve their structure with cryo-EM. We provide biochemical and structural evidence that NLRP1-CARD and CARD8-CARD filaments have different intrinsic affinities for the adaptor ASC. Despite very similar helical parameters, the subtle differences in rotational angles of individual NLRP1-CARD and CARD8-CARD monomers and their singular Type II junctions give rise to distinct surface charge distributions at the filament scale. Extending this analysis, we also report the specific Type II interface motifs that directly dictate the compatibility of heterotypic pairwise CARD-CARD interactions among NLRP1, CARD8, NLRC4, ASC and CASP1. Taken together, these data demonstrate that the CARD domains are not mere interchangeable ‘lego building blocks” for large prionoid oligomers. Instead, each possesses intrinsic abilities to discriminate amongst potential downstream binding partners in order to specify the composition of different inflammasome complexes. It is conceivable that these distinct inflammasome complexes can lead to distinct biological outcomes, in agreement with previous findings demonstrating that different strengths of the pyroptotic signaling are required for cell death and IL1 cytokine secretion ^36,37^. Interestingly, the prominent role in Type II interface-dependent assembly is apparently specific to the human inflammasome complexes, as other CARD-domain innate immune complexes, such as MAVS (EMD-5925)^28^, BCL10 (EMD-7314)^38^, RIPK2 (EMD-4399, 6842)^30,39^ rely more heavily on Type I interface junctions (e.g., MAVS Arg43-Typ56 density bridge). This underscores the importance of studying molecular structures of large complexes at atomic resolutions. As supra-molecular assemblies are highly prevalent in innate immune and cell death signaling ^40^, our findings should inform future work to decipher the general assembly principles of key molecular complexes involved in these processes.

## Acknowledgments

We thank all members of the Zhong lab, Reversade lab and Wu lab for their support and help. We acknowledge useful discussions with Dr. Luo Dahai (NTU, Singapore) and Dr. Seth Masters (WEHI, Australia). This work is supported by the Agency for Science, Technology and Research (GODAFIT Strategic Positioning Fund, Bruno Reversade), National Medical Research Council, Singapore (NMRC/OFYIRG/0046/2017, Franklin Zhong), Concern Foundation’s Conquer Cancer Now Award (Franklin Zhong), National Research Foundation, Singapore (Franklin Zhong), Nanyang Assistant Professorship (Wu Bin, Franklin Zhong), MOH-NMRC-OFIRG grant (MOH-000382-00) and Tier II Ministry of Education Research Grant (MOE2016-T2-1-010-Wu Bin). Bruno Reversade is a fellow of the Branco Weiss Foundation and a recipient of the A*STAR Investigatorship, Senior NRF investigatorship and EMBO Young Investigatorship. Franklin Zhong and Wu Bin are Nanyang Assistant Professors. Franklin Zhong is a recipient of the National Research Foundation Fellowship.

## Author contributions

Gong Qin prepared NLRP1 protein samples, conducted all *in vitro* biochemical experiments, prepared EM samples, collected EM data, and analyzed the results. Kim Robinson and Daniel Teo carried out all cellular experiments and microscopy. Xu Chenrui prepared NLRC4 protein samples, conducted imaging processing and computational work. Zhang Jiawen prepared a CARD8 sample and conducted computational work. Boo Zhao Zhi prepared ASC proteins, conducted biochemical assays related to ASC. Zhang Yaming facilitated all protein expression and participated in cloning work. Kim Robinson, John Lim, Goh Wah Ing and Graham Wright jointly conducted 3D-SIM experiments and data analysis. Wu Bin performed cryo-EM data processing. Franklin Zhong, Wu Bin and Bruno Reversade jointly conceived of the project, designed and coordinated the experiments, analyzed the results and wrote the manuscript.

## Methods

### Cell culture

HEK293Ts (ATCC #CRL-3216) were obtained from commercial sources and cultured according to the suppliers’ protocols. Immortalized human keratinocytes (N/TERT-1) were a kind gift from H. Reinwald (MTA) ^41^. All cell lines underwent routine Mycoplasma testing with Lonza MycoAlert (Lonza #LT07-118).

### Transient transfection and stable cell line generation using lentiviruses

293T-ASC-GFP cells were described previously ^8^. All other 293T-GFP responder cell lines were constructed by lentiviral transduction (pCDH vector backbone, System Biosciences). All transient expression plasmids were cloned into the pCS2+ vector using standard restriction cloning. Polyclonal Cas9/CRISPR knockout cell lines were generated with lentiCRISPR-v2 (Addgene #52961) and selected with puromycin. Knockout efficiency was tested with Western blot 7-10 days after puromycin selection. Site-directed mutagenesis was carried out with QuickChangeXL II (Agilent #200522).

### 3D structured illumination microscopy (3D-SIM) & widefield-deconvolution

A DeltaVision OMX v4 Blaze microscope (GE Healthcare) equipped with 488nm and 568nm lasers for excitation (3D-SIM) and solid state illuminator (widefield) the BGR-FR filter drawer (emission wavelengths 436/31 for DAPI) was used for acquisition of 3D-SIM and widefield-deconvolution images. An Olympus Plan Apochromat 100x/1.4 PSF oil immersion objective lens was used with liquid-cooled Photometrics Evolve EM-CCD cameras for each channel. For 3D-SIM, 15 images per section per channel were acquired (made up of 3 rotations and 5 phase movements of the diffraction grating) at a z-spacing of 0.125 µm. The same z-spacing was used for widefield acquisition. Structured illumination reconstruction or deconvolution and alignment was completed using the SoftWorX (GE Healthcare) program, 3D image rendering and analysis in Imaris version 9.3.0 (Bitplane an Oxford Instruments Company), 2D image analysis in Fiji (Schindelin et al., 2012) and figure preparation in Illustrator.

### Antibodies and cytokine analysis

The following antibodies were used in this study: HA tag (Santa Cruz Biotechnology, #sc-805), GAPDH (Santa Cruz Biotechnology, #sc-47724), ASC (Adipogen, #AL-177), CASP1 (Santa Cruz Biotechnology, #sc-622), IL1B (R&D systems, #AF-201), FLAG (SigmaAldrich, #F3165), GFP (Abcam, #ab290). IL-1B measurements were carried out with human IL-1B ELISA kit (BD, #557953).

### Plasmid construction for recombinant protein expression

For recombinant CARD proteins, NLRP1 constructs (985-1473, 1213-1473, 1374-1466, 1374–1473) with or without C-terminal SNAP tag (NEB), CARD8–CARD (451-537) with or without C-terminal SNAP tag, ASC–CARD (1–102) with N-terminal SNAP tag, CASP1-CARD (1-92) with or without C-terminal SNAP tag, NLRC4-CARD (1-102) with or without C-terminal SNAP tag. were cloned into pET47 vector in between XmaI and XhoI restriction sites. A 3C protease cleavage site was inserted in between the CARD domain and the SNAP tags in case SNAP needs to be removed for biochemical analysis. Site-directed mutagenesis was performed using the KAPA Hifi PCR kits (KAPA Biosystems).

### Protein expression and purification

For the expression of the His-tagged recombinant proteins, the constructed plasmids were transformed into BL21(DE3) (NEB) and grown at 37□°C for 16–18□h. Then the starting culture was transferred to a larger volume of lysogeny broth (LB), which was grown until an OD600 of about 0.6–0.8, then the temperature was lowered to 16□°C, and the cells were induced with 0.5□mM IPTG and grown overnight. Bacteria were harvested at 4600□×□g (Avanti JXN series, Beckman Coulter) for 10□min and then resuspended with lysis buffer (20□mM Tris-HCl, pH 8.0, 300□mM NaCl, 20□mM imidazole, and 10% glycerol). Cells were then lysed by Emulsiflex-C3, and centrifuged at a speed of 30,000□×□g (Avanti JXN series, Beckman Coulter) for 20□min at 4□°C to separate supernatant and pellet. The supernatant was passed through a pre-equilibrated column containing Ni-NTA agarose beads (ThermoFisher Scientific) by gravity. The column was then washed with lysis buffer containing 20□mM imidazole to remove nonspecific binding proteins. His-tagged proteins were eluted with elution buffer that contained 300□mM imidazole. To further purify our recombinant proteins, size-exclusion chromatography (Superdex 200 increase, GE Healthcare) was performed in buffer A (20□mM Tris-HCl, pH 8.0, 150□mM NaCl, and 1□mM DTT). At this step, the purified proteins were the oligomeric pre-activated seed fraction. For monomeric CARD-SNAP or SNAP-CARD domain proteins, it was prepared following the protocol as described here. Concentrated and purified soluble filament seed at around 10□mg/ml was buffer exchanged into 10□ml of 6□M guanidinium, and gradually dialyzed against 3, 2, 1.5, 1, 0.8, and 0.6□M guanidinium containing buffer A in the cold room, and eventually in buffer A. In the absence of stimuli, refolded SNAP-ASC-CARD protein remained monomeric for 24□h, and CASP1-CARD-SNAP remained monomeric or lower oligomeric for about 12 h. Recombinant SNAP-NLRP1-FIIND^UPA^-CARD and SNAP-CARD8-FIIND^UPA^-CARD were expressed in similar ways as the CARD domain constructs. Notably, most of the protein exist in cellular pellets after the Emulsiflex step, thus, direct pellet wash with lysis buffer with additional 1% Triton-X100, followed by 6 M Guanidinium is necessary to obtain solution proteins. Chemical refolding following the same dialysis protocol, followed by 3C cleavage of the SNAP tag, was used to obtain homogenous oligomeric population of FIIND^UPA^-CARD complexes at various concentrations.

### *In vitro* Oligomerization assay

SNAP-ASC-CARD and CASP1-CARD-SNAP monomers that were labeled with Alexa Fluor 647 (New England Biolabs) were co-incubated with CARD domain only filament seeds (CARD8-CARD without SNAP, NLRP1-CARD without SNAP, NLRC4-CARD without SNAP) separately. One microliter of 100□μM BG-Alexa dye stocks (dissolved in water containing 5% DMSO) were added into every 100□μl of pre-labeling protein stock at a concentration of 20□μM. Fluorescently labeled monomeric SNAP-ASC-CARD (10□μM) and CASP1-CARD-SNAP (10□μM) was mixed with 20 or 5 μM seed oligomer for 15□min at room temperature before analysis by Bis-Tris native PAGE gel (Life) or 15% SDS-PAGE gel. The fluorescently labeled gel was run at a voltage of 200□V for 60□min. Once the EMSA was completed, they were visualized by Typhoon FLA7000 (GE Healthcare) with 100-μm pixel size at channel 647 using PTM 600. To form ASC-CARD and NLRC4-CARD long filament (200–500□nm), ∼20□μM monomeric SNAP tagged protein was mixed with unlabeled, wild-type SNAP tagged ASC-CARD and NLRC4-CARD seeds at a molar ratio of 20:1 for overnight. Later on, in-house-prepared MBP-3C-protease was added to the fully extended filament with a final concentration of 0.5□μM. The digestion was performed at 4° for 8□h, before further size exclusion purification to separate the core filament, the cleaved-off SNAP tags, and proteases. Final extended ASC–CARD and NLRC4-CARD thin filament were then purified using size-exclusion column Superose 6 (GE Healthcare), before structural analysis. To form long NLRP1-CARD and CARD8-CARD filaments, N-terminal His-tagged CARD domain alone constructs were expressed and purified according to the protocol mentioned above, then concentrated to around 400 μM, homogeneous long filaments formed spontaneously, without the requirement of seeding.

### Negative stain-electron microscopy

Samples were applied onto the grids and stained following a Harvard medical school Liao lab protocol (https://liao.hms.harvard.edu/node/32, originally designed by Tom Walz). Protein samples were diluted to 0.01 to 0.05 mg/ml for best sample density on the grids, immediately before grid preparation to prevent concentration dependant disassociation. Images were taken using a T12 120keV TEM microscope, at either 30,000X or 48,000X magnification, and calibrated against known objects of MAVS-CARD filament.

NLRP1-FIIND^UPA^-CARD complex density model was built based on 58 negative stained EM images with homogenous background, total 818 segments from a single 15 nm thick 2D class. Combining more segments yielded very noisy maps. Several preliminary maps ∼29 Å were obtained after refinement, when different helical parameters were adopted, but all maps have the similar two layer features. In addition, such strong inner layer density and spotted outer layer density are always seen in all solutions. The illustrated 3D map was obtained when choosing a helical parameters similar to NLRC4 density EMDB-2901 and analysis of the NLRP1-FIIND^UPA^-CARD 2D class images. We were cautious about the accuracy of this particular map, thus simply using it as a guide to piece together an overall architecture of NLRP1 inflammasome.

### Cryo-electron microscopy

For cryo-EM grid preparation, Quantifoil R1.2/1.3 holey grid (Quantifoil) was glow-discharged for 60s and plunge-frozen using a Vitrobot (FEI) to be treated with 5-µl sample at a concentration of about 2 mg/ml. Final high-resolution images were collected at Titan Krios, 300□kV, using Gatan K2 camera (Gatan), super-resolution mode for data collection, and binned 2× when doing motion correction. The sub-frame time is 250□ms and total exposure time is 10□s, this will give 40 frames per stack. The pixel size on the final image is 1.10□Å. The dose rate is 4□e/pixel/s.

For cryo-EM data, super-resolution image stacks were gain normalized and binned by 2× to a pixel size of 1.10□Å prior to drift and local movement correction using MotionCorr2. Date processing software Relion 3.0 was used for CTF determination, particle picking, 2D classification, 3D classification, and refinement procedures. Totally, 2187 (NLRP1), 1424 (CARD8), 3183 (ASC), and 2109 (NLRC4) micrographs were collected and analyzed. We manually picked out ∼ 10,000 to 20,000 filaments for each sample and windowed out segments of 240□×□240 pixels, yielding 422,388 (NLRP1), 1,201,108 (CARD8), 877,406 (ASC-CARD) and 382,715 (NLRC4) particles for 2D classification. After several iterations, bad particles and less desirable classes were removed and we used 100,240 (NLRP1), 301,848 (CARD8), 131,260 (ASC-CARD) and 120,602 (NLRC4) particles for 3D classification. Ab initio low-resolution helical structure was generated using a Gaussian cylinder as an initial model. 3D classification and auto-refinement of 53,241 (NLRP1), 98,280 (CARD8), 59,566 (ASC-CARD) and 120,602 (NLRC4) particles were done in Relion3.0 and CisTEM. All 3D refinements were carried out following the gold-standard procedure where the data set was divided into two half-sets. After refinement was converged, a mask of the central 30% segment was calculated and applied to the final data set and the corrected Fourier shell correlation (FSC) was calculated to estimate the resolution about 3.7□Å (NLRP1), 3.7□Å (CARD8), 4.1□Å (ASC-CARD), 3.3□Å (NLRC4) by using FSC□=□0.143 criterion. In the final model, parameters of the RIP2 filament were calculated to yield 5.36□Å (NLRP1), 5.409□Å (CARD8), 5.27□Å (ASC-CARD) and 5.17□Å (NLRC4) of the axial rise per asymmetric unit and an azimuthal rotation per subunit of −100,821° (NLRP1), −99,16° (CARD8), −100,614° (ASC-CARD), −100,55° (NLRC4).

### Model building

Unpolished maps were used for model building.

Step 1 was the monomer docking. The CARD monomer structures (PDB-4IFP for NLRP1-CARD, PDB-4KIM for CARD8-CARD, PDB-6N1H for ASC-CARD, and PDB-6N1I for NLRC4-CARD) were used as starting points at this stage. In the cases of NLRP1 and CARD8, the residue numbers were renumbered and the terminal residues were deleted to match the length of the density. The individual helix was manually shifted (rigid body movement) according to the locations of three large aromatic side chains. The density map at this resolution enabled us to unambiguously locate these residues. This step was done in Coot manually.

In Step 2, a tetramer containing the Type I, II, and III interfaces (a centrally located monomer and three other monomers forming Type I, II, and III interfaces with the central one) was built. After building four monomers in this way, we obtained a tetramer that could be used to refine the surface interactions. Phenix real space refinement was done without rigid body refinement. Different parameters were tested, a torsion-rotational angle NCS constraints (Phenix default auto setting) was used at this step. In this way, all surfaces of the central monomer could be refined.

During Step 3, a tetramer model was built that is in agreement with the data from the cellular activity assays, given particular emphasis in charge-reversal experiments. In Step 4, a 12mer model was built. The central monomer from Step 3 was copy-and-pasted to occupy the entire 60□Å segment density. In the end, 12 identical monomers were added to the density. Phenix real space refinement with default NCS torsion-rotational constraint was performed. After several rounds of rigid body refinement, a model with reasonable statistics was obtained.

Step 5: final refinement. Specific issues, like high clash score and Ramachandran plot outliers after Step 4 were manually identified and rebuilt and then Step 4 was repeated until satisfactory statistics and density interpretation were achieved.

### Data Availability Statement

The densities and structure models discussed in this study were validated and deposited in public databases. NLRC4-CARD filaments (PDB-6K8J, EMD-9946), ASC-CARD filaments (PDB-6K99, EMD-9947), NLRP1-CARD (PDB-6K7V, EMD-9943), CARD8-CARD (PDB-6K9F, EMD-9948)

## Legends for Extended Data Figure S1-S8

**Fig. S1.**
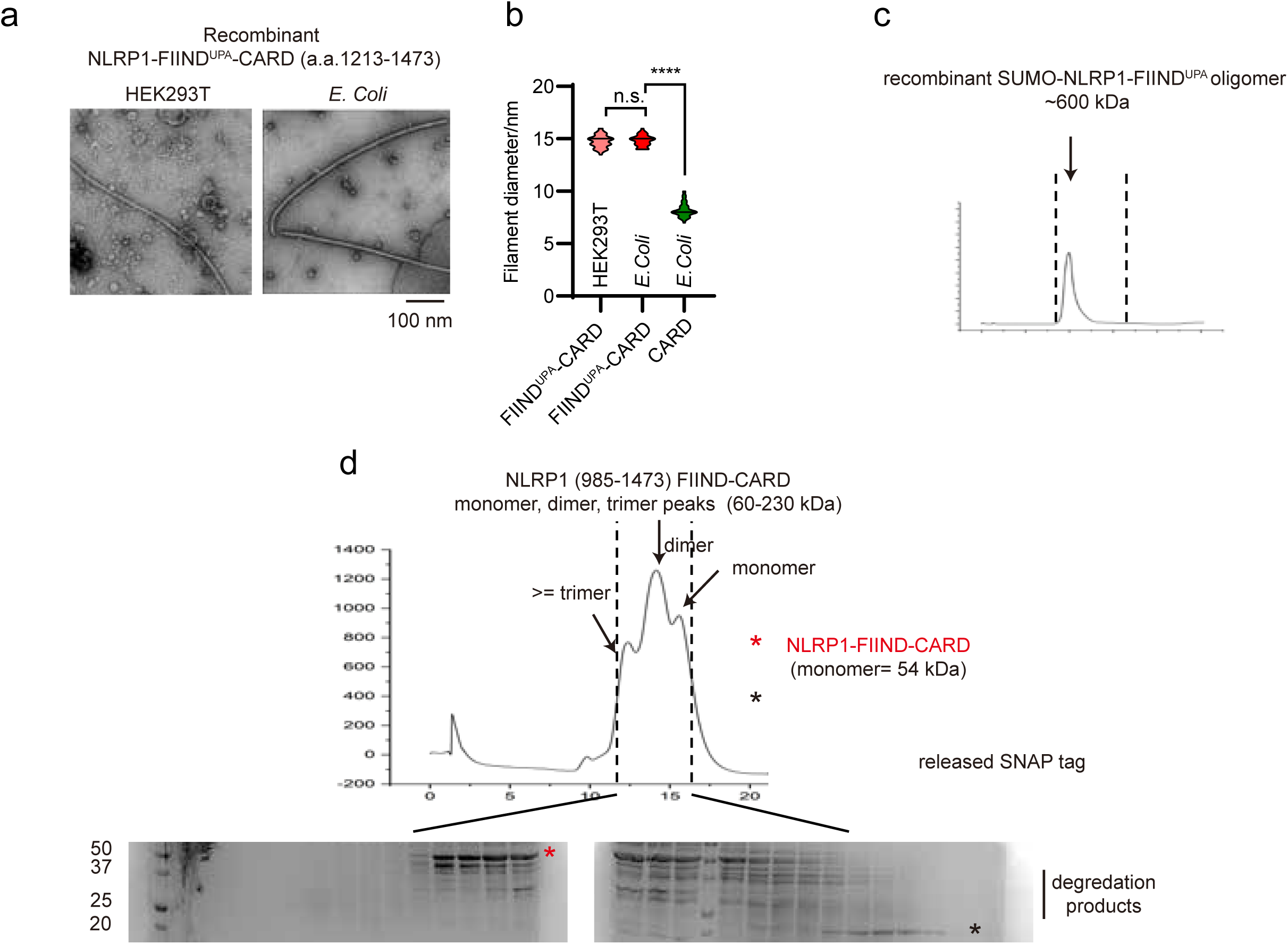
Additional data for the oligomerization of NLRP1 constructs. a. Negative EM images of NLRP1-FIND^UPA^-CARD filaments purified from HEK293T cells and *E*.*Coli*. b. Dimensions of NLRP1-FIIND^UPA^-CARD and NLRP1-CARD filaments, based on negative stain EM images, ∼100 individual measurements. P-value was calculated with One-way ANOVA. c. Size exclusion profile of purified SUMO-NLRP1-FIIND^UPA^ on Superdex S200 increase column. d. Size exclusion profile and SDS-PAGE analysis of NLRP1-FIIND^fl^-CARD (a.a. 985-1473) construct. Unlike FIIND^UPA^-CARD, FIIND^fl^-CARD appears to be a monomer-dimer mixture, instead of a 15-20 mer mixture.

**Fig. S2.**
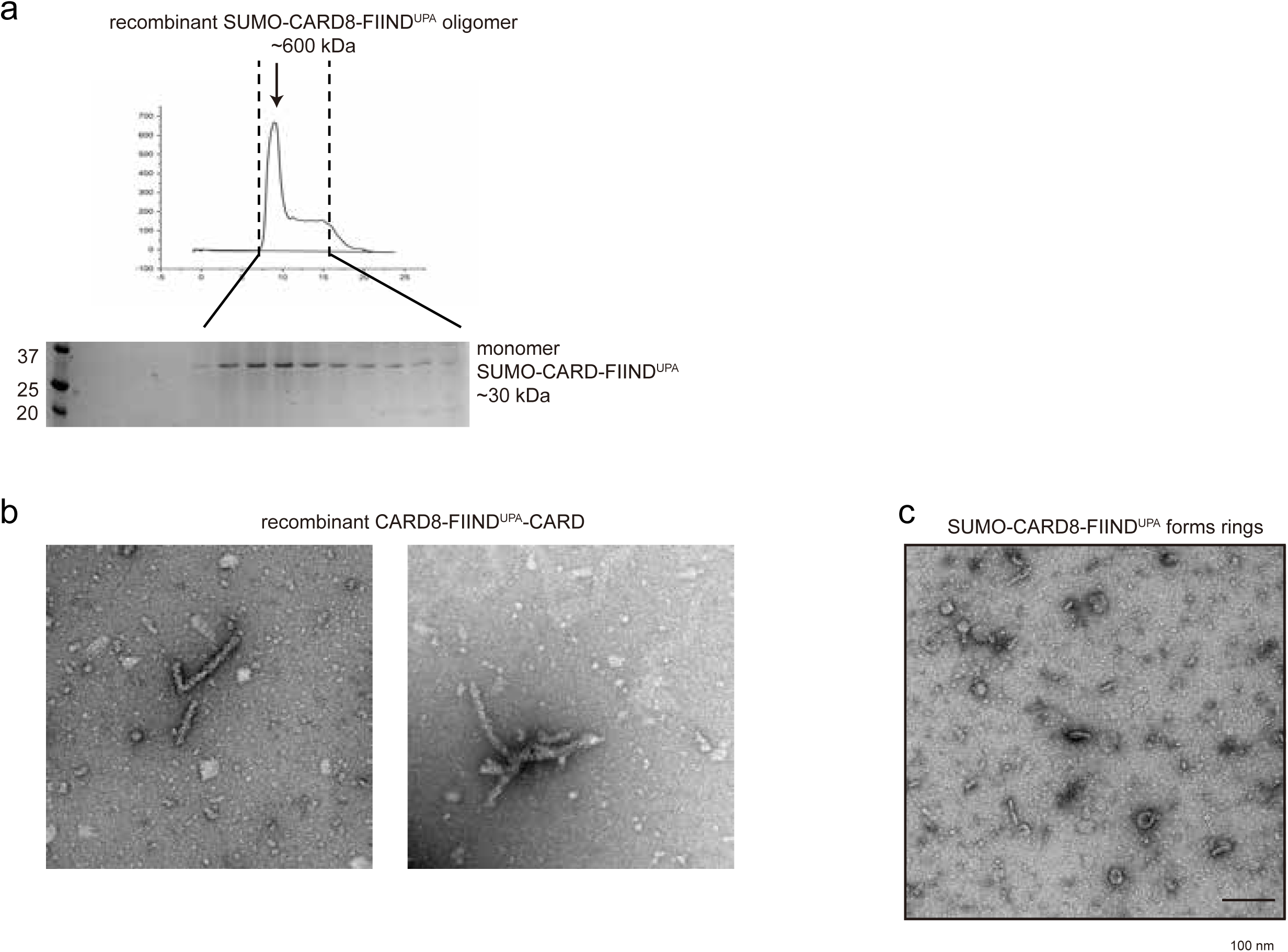
Additional data for the oligomerization of CARD8 constructs. a. Size exclusion and SDS-PAGE profile of purified SUMO-CARD8-FIIND^UPA^, Superdex S200 increase column. b. Negative EM images of CARD8-FIND^UPA^-CARD complexes purified from *E*.*Coli*. c. Negative EM images of CARD8-FIND^UPA^ ring-complexes purified from *E*.*Coli*.

**Fig. S3.**
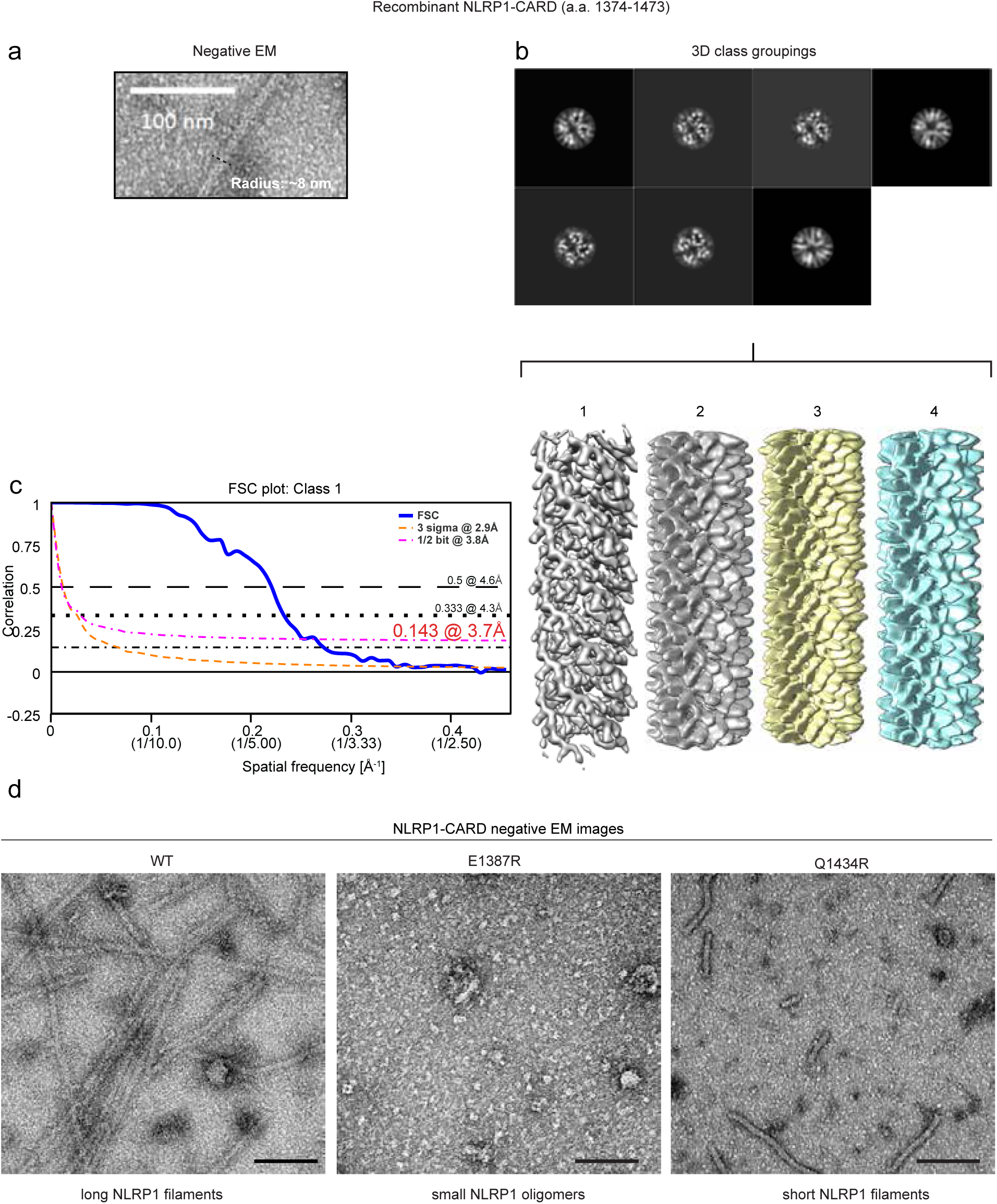
Additional data for NLRP1-CARD filament analysis. a. Representative dimensions of NLRP1-CARD filaments. b. 3D densities of 4 groups of NLRP1-CARD filaments after helical symmetry was applied. c. FSC plot of the final post-refinement NLRP1-CARD filament density (EMD-9943). d. Negative stain EM images of recombinant wild-type NLRP1-CARD, NLRP1-CARDE^1387R^ and NLRP1-CARD^Q1434R^.

**Fig. S4.**
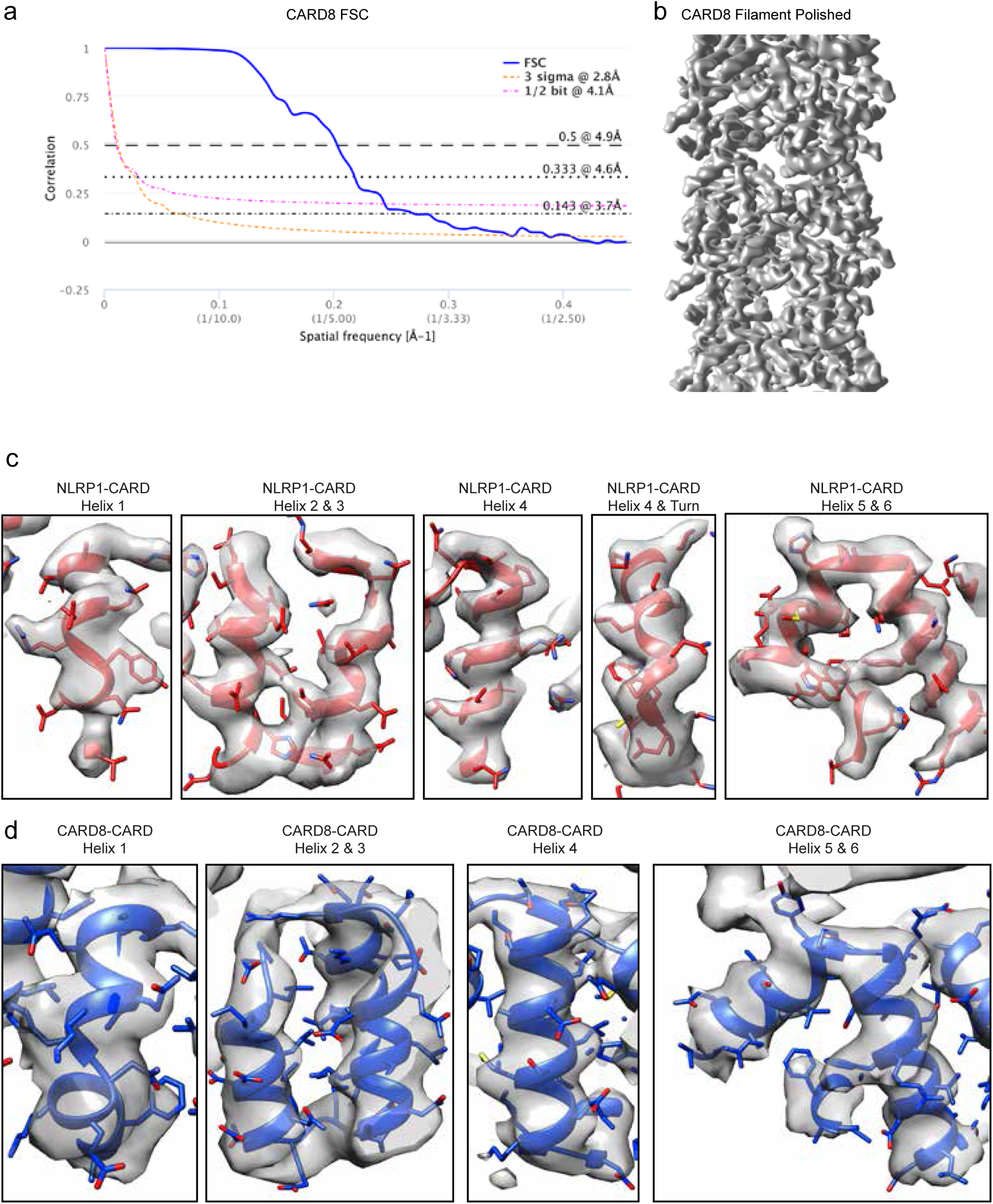
Additional data for NLRP1-CARD and CARD8-CARD complex densities. a. FSC plot of final refined CARD8-CARD filament density (EMD-9948). b. Post-refinement 3D density of CARD8-CARD filaments. c. Assignment of individual residues and α-helices in NLRP1-CARD structure. d. Assignment of individual residues and α-helices in CARD8-CARD structure.

**Fig. S5.**
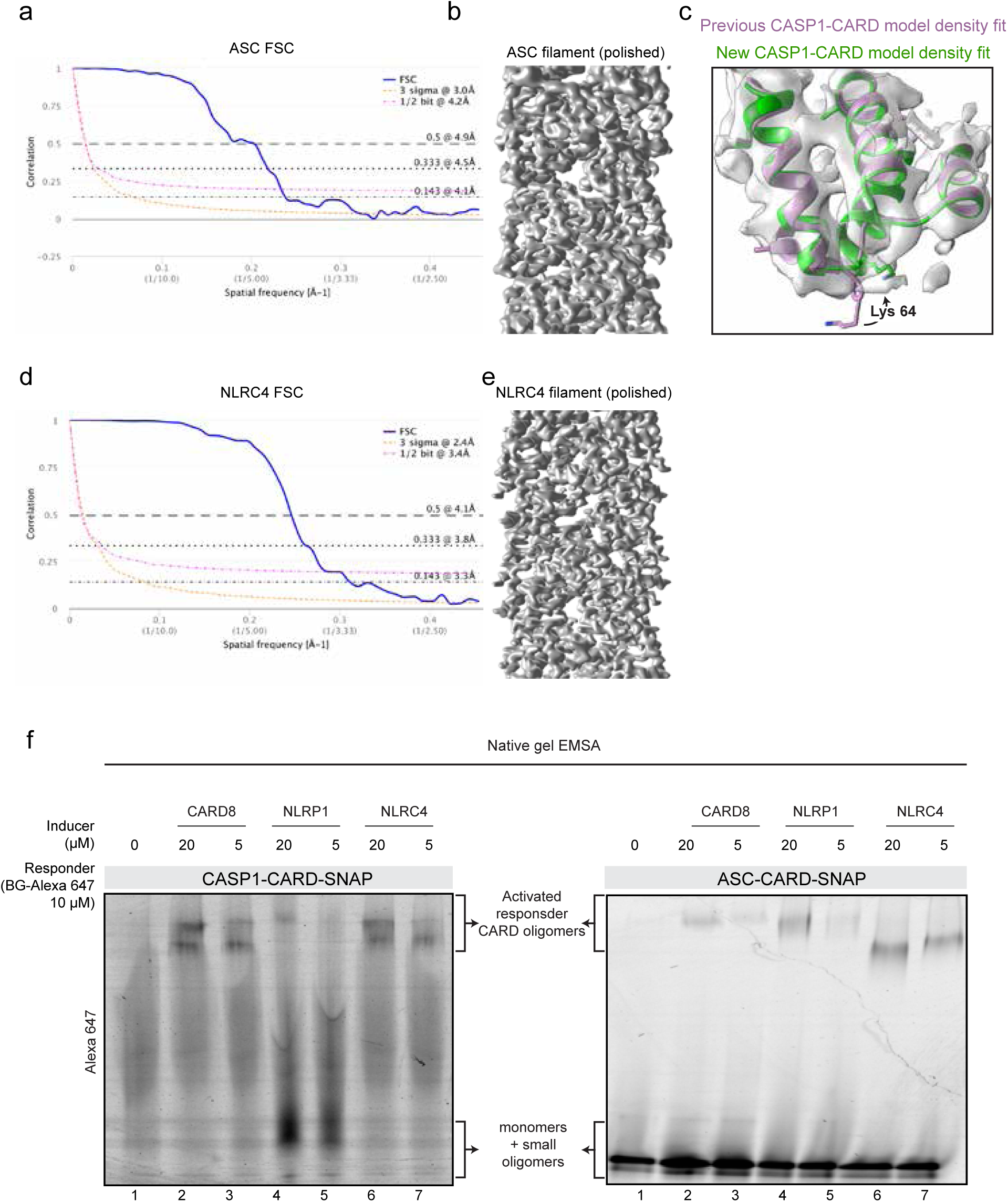
Additional data for ASC-CARD, CASP1-CARD and NLRC4-CARD structural analysis. a. FSC plot of final, post-refinement ASC-CARD filament density (EMD-9947). b. Post-refinement 3D density of ASC-CARD filament. c. Refined fitting of CASP1-CARD density (EMD-3241). Purple is original PDB fitting (PDB-5FNA). Green is our new, refined fitting based on the newly acquired, homologous CARD8-CARD structure. A shift of amino acid register was identified, supported by Lys64 density assignment. d. FSC plot of final, post-refinement NLRC4-CARD filament density (EMD-9946). e. Post-refinement 3D density of NLRC4-CARD filament. f. Left: native gel shift assay of recombinant CASP1-CARD-SNAP oligomerization induced by purified CARD8-CARD, NLRP1-CARD and NLRC4-CARD. Right: native gel shift assay of ASC-CARD-SNAP oligomerization induced by purified CARD8-CARD, NLRP1-CARD and NLRC4-CARD.

**Fig. S6.**
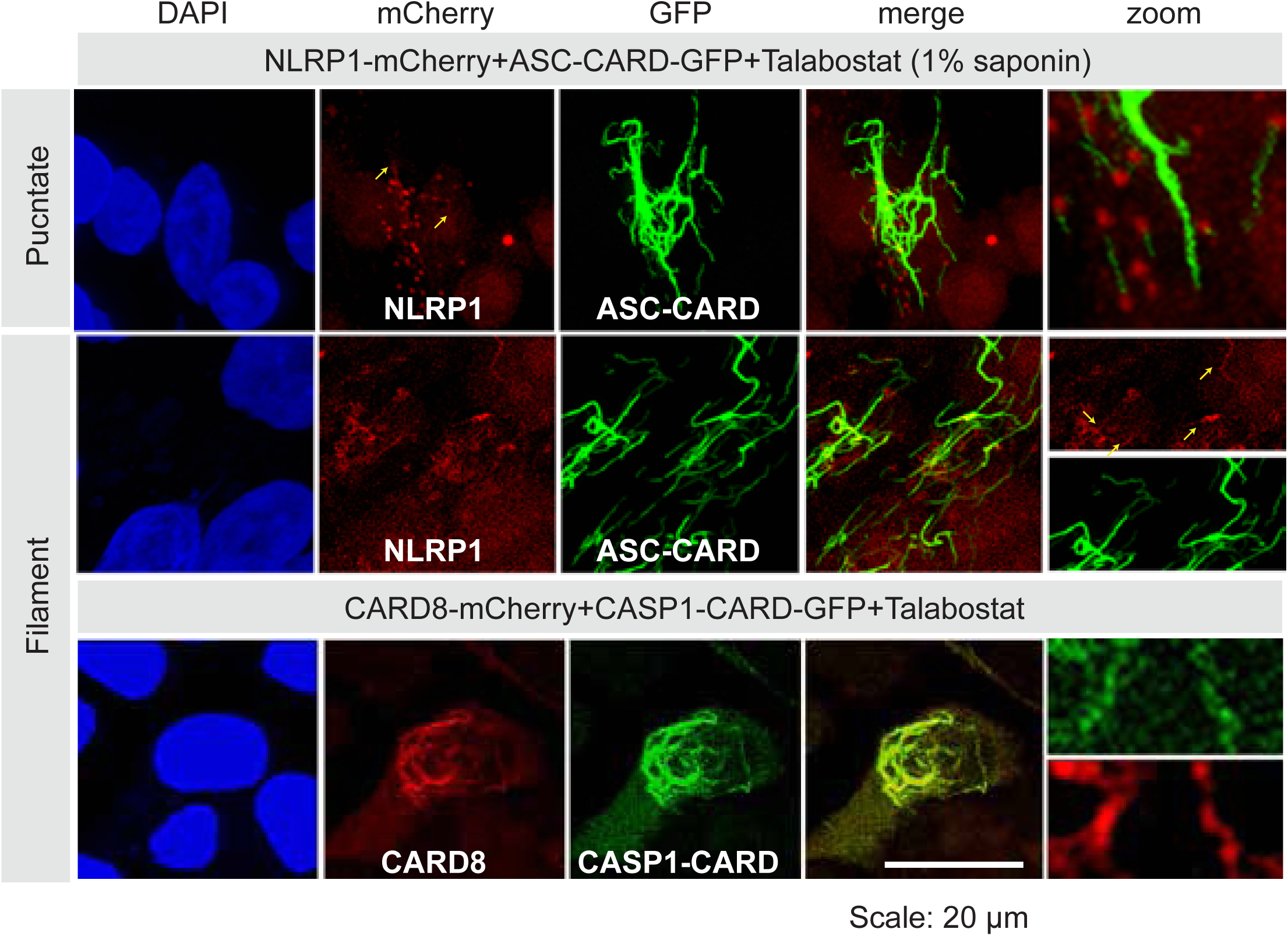
Full-length NLRP1- and CARD-mCherry filaments observed in Talabostat-treated 293T cells.

**Fig. S7.**
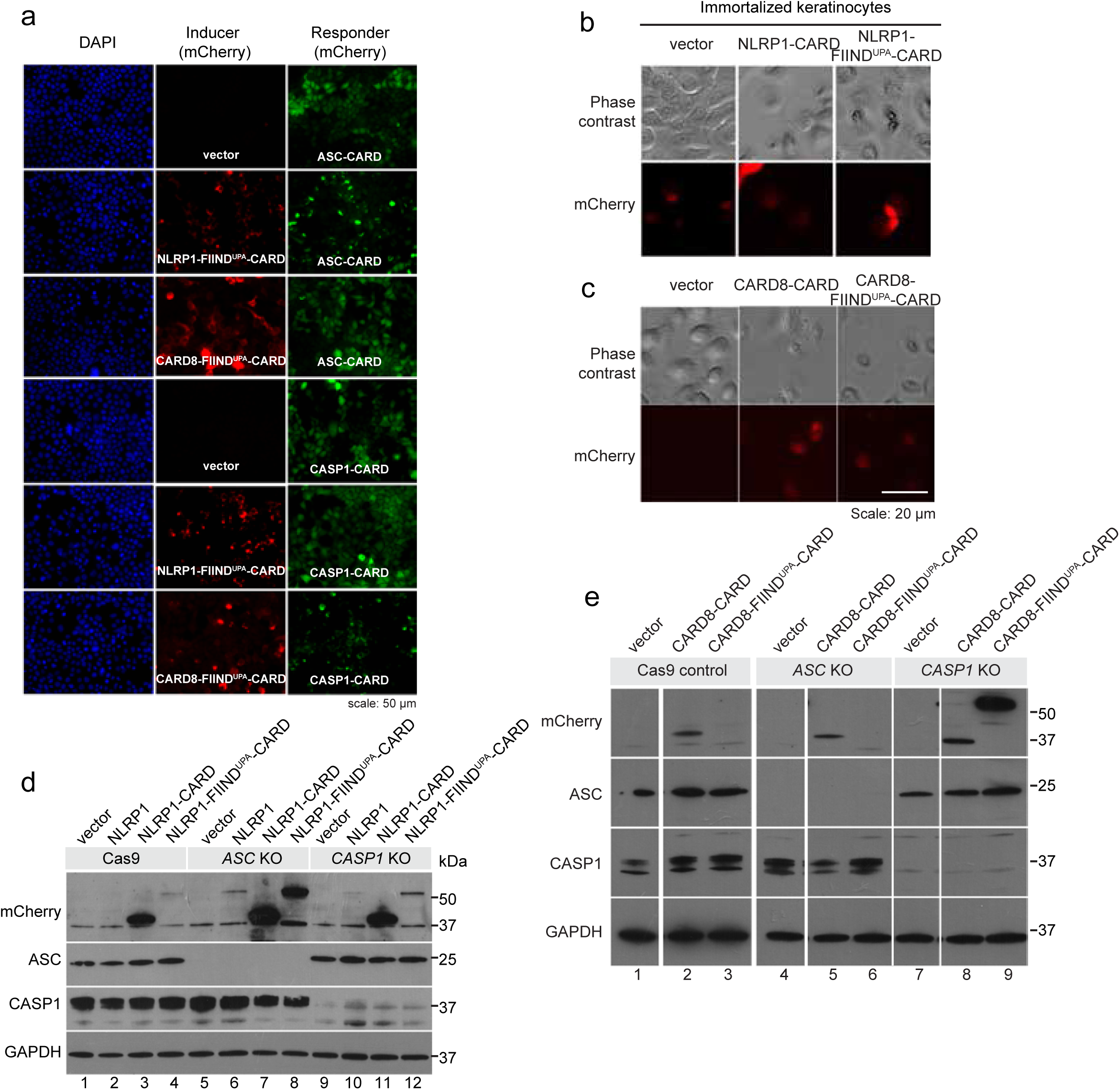
Support data of NLRP1 and CARD8 inflammasome assembly in mammalian cells. a. Representative images of 293T-ASC-CARD- and CASP1-CARD-GFP responder cells transfected with mCherry-tagged NLRP1/CARD-FIIND^UPA^-CARD or NLRP1/CARD8-CARD. b. Representative images of wild-type immortalized human keratinocytes transfected with mCherry-tagged NLRP1-CARD, NLRP1-FIND^UPA^-CARD. c. Representative images of wild-type immortalized human keratinocytes transfected with mCherry-tagged CARD8-CARD and CARD8-FIND^UPA^-CARD. d. Western blot of mCherry-tagged NLRP1 constructs in control, *ASC* and *CASP1* KO keratinocytes. e. Western blot of mCherry-tagged CARD8 constructs in control, *ASC* and *CASP1* KO keratinocytes.

**Fig. S8.**
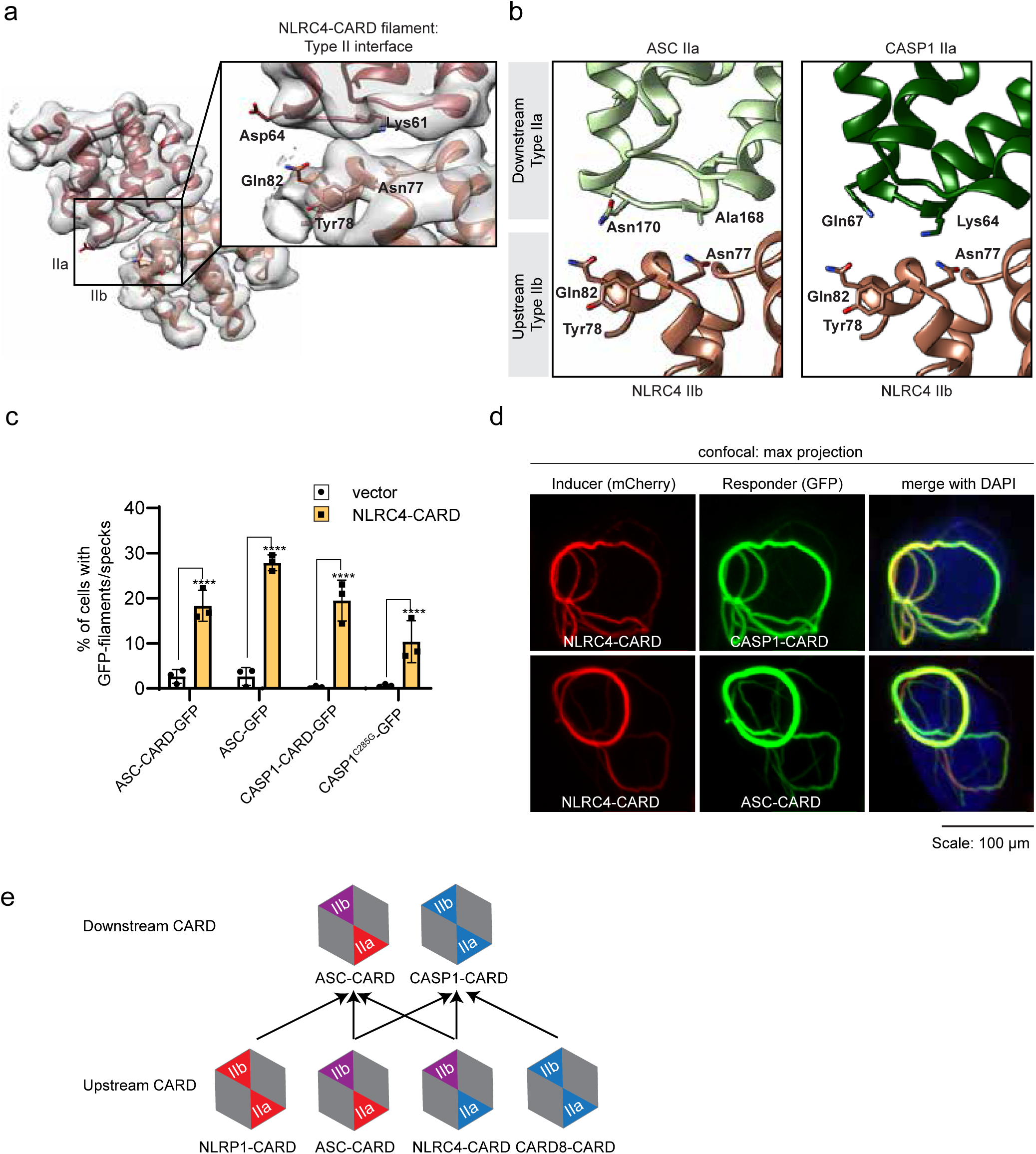
Proposed structural interpretation of NLRC4 inflammasome assembly. a. Atomic model and directly observed density map at Type II interface of human NLRC4-CARD filaments (PDB-6K8J, EMD-9946). b. Type II junction interactions between NLRC4-CARD and ASC-CARD (left), and between NLRC4-CARD and CASP1-CARD (right). c. Percentage of filament/speck formation in 293T-ASC-CARD-GFP, ASC-GFP, CASP1-CARD-GFP and CASP1C^285G^-GFP responder cells transfected with NLRC4-CARD-mCherry. P-value was calculated with One-way ANOVA, n=3 transfections. d. Confocal microscopy images of NLRC4-mCherry filaments with CASP1-CARD-GFP and ASC-CARD-GFP filaments in 293T cells. e. Summary of unidirectional CARD-CARD interactions in human NLRP1, CARD8 and NLRC4 inflammasome complexes.

