## Supplemental Table 1 for "Structural basis for distinct inflammasome complex assembly by human NLRP1 and CARD8"

Supplement Table 1. Data Collection, map and model refinement, validation

| | ASC-CARD (PDB-6K99, EMD-9947) | NLRC4-CARD (PDB-6K8J, EMD-9946), | CARD8-CARD (PDB-6K9F, EMDB-9948) | NLRP1-CARD (PDB-6K7V, EMDB-9943) |
|---|---|---|---|---|
| **Data Collection** | | | | |
| Microscope | Titan Krios | Titan Krios | Titan Krios | Titan Krios |
| Voltage (kV) | 300 | 300 | 300 | 300 |
| Detector | K2 | K2 | K2 | K2 |
| Pixel size (Å) | 1.1 | 1.1 | 1.1 | 1.1 |
| Defocus range (µm) | -1,-2 | -1,-2 | -1,-2 | -1,-2 |
| Electron Dose ($e^-/A^2$) | 40 | 40 | 40 | 40 |
| **Helical Reconstruction** | | | | |
| Software | RELION 3.0beta | RELION 3.0beta | RELION 3.0beta | RELION 3.0beta |
| Segment length (Å) | 220 | 220 | 264 | 264 |
| Particles | 877408 | 382715 | 1201108 | 422388 |
| Helical rise (Å) | 5.27 | 5.17 | 5.409 | 5.36 |
| Helical rotation (°) | -100.614 | -100.55 | -99.16 | -100.821 |
| Resolution (Å) | 4.1 | 3.3 | 3.7 | 3.7 |
| **Coordinate Refinement (mid-segment 12mer)** | | | | |
| Software | Phenix | Phenix | Phenix | Phenix |
| Rwork | 0.211 | 0.2571 | 0.2530 | 0.2548 |
| Rfree | 0.2208 | 0.3057 | 0.3017 | 0.2846 |
| **Model** | | | | |
| Number of residues | 996 | 1020 | 1044 | 1008 |
| B-factor overall | 206 | 249 | 276 | 360 |
| R.M.S. deviation | | | | |
| Bond length (Å) | 0.003 | 0.002 | 0.002 | 0.005 |
| Bond angle (°) | 0.670 | 0.364 | 0.423 | 0.767 |
| **Validation** | | | | |
| Molprobity clashscore | 6.92 | 0.24 | 5.41 | 2.06 |
| Rotamer Outliers (%) | 1.3 | 0 | 0 | 1.5 |
| $C_\beta$ deviation (%) | 0 | 0 | 0 | 0 |
| **Ramachandran plot** | | | | |
| Fovored (%) | 92.5 | 98.1 | 100 | 96.5 |
| Allowed (%) | 6.2 | 1.9 | 0 | 3.5 |
| Outliers (%) | 0 | 0 | 0 | 0 |
